# The Landscape of tRNA Modifications in Archaea

**DOI:** 10.1101/2025.05.02.651894

**Authors:** Jesse S. Leavitt, Henry Moore, Thomas J. Santangelo, Todd M. Lowe

## Abstract

Transfer RNA (tRNA) modifications are essential for structural integrity, decoding fidelity, and stress adaptation, yet their evolutionary dynamics remain poorly understood. Here, we apply Ordered Two-Template Relay sequencing (OTTR-seq) to comprehensively profile tRNA modifications across nine archaeal species spanning diverse ecological niches. We uncover coordinated and mutually exclusive methylation at acceptor stem positions 6 and 67 in hyperthermophiles, as well as clade-specific co-modification at positions 10 and 26, which are typically known for their importance as tRNA modification anti-determinants. Comparative analyses also reveal lineage-specific divergence in the domain architectures of tRNA methyltransferases, including Trm14, Trm10, Trm11, and Trm1. We further refine known identity elements such as the G10oU25 pairing, and highlight novel structural contexts that facilitate or prevent modification. These findings exemplify the co-evolution of tRNAs and their modifying enzymes, providing new insights into how archaea may fine-tune translation in extreme environments. The scope of these data and comparative analyses establish a multispecies framework for future biochemical, mechanistic, and predictive modeling efforts.

## Introduction

Transfer RNAs (tRNAs) play a central role in translating genetic information into functional proteins in all living systems. Post-transcriptional modifications heavily influence tRNA structure and function, modulating the rate of maturation, molecular stability, and the specificity of interactions with aminoacyl-tRNA synthetases, mRNA codons, and a multitude of other cellular components (Grosjean et al., 2008; Grosjean, 2010; Hernandez-Alias et al., 2022; Schultz & Kothe, 2024; Sokołowski et al., 2018; Yared et al., 2024). These modifications are differentially conserved at distinct positions within tRNAs, imparting specific properties that vary considerably across the tree of life (Höfer & Jäschke, 2018; Ontiveros et al., 2019).

Archaea, as the least-studied domain, offers unique insights into how RNA-based regulation and stability evolved, highlighting both the ancient strategies that enabled translation under extreme conditions and the evolutionary roots of RNA modification systems in eukaryotes. Archaea’s ecological niches include temperatures approaching 100°C in deep-sea hydrothermal vents (Schicho et al., 1993), acid hot springs with pH values near 2 (Reno et al., 2009), and hypersaline environments with salt concentrations exceeding 3M NaCl (Gupta, 1984). Among methanogens, found exclusively within Archaea, adaptation to anoxic and high-pressure marine environments is evident (Whitman et al., 1987).

For this fascinating branch of life, the complete patterns of tRNA modifications are known for only two species, primarily due to the high cost, difficulty, and low-throughput nature of traditional biochemical characterization of tRNAs. From the comprehensive mapping in *Haloferax volcanii* and *Methanocaldococcus jannaschii* (Gupta, 1984; Yu et al., 2019), along with partial mapping in *Methanococcus maripaludis*, *Pyrococcus furiosus*, and *Sulfolobus acidocaldarius* (Wolff et al., 2023), it is clear that tRNA modifications contribute to adaptation in diverse extreme environments (Fluke et al., 2024; Hori et al., 2018; Kowalak et al., 1994; Li et al., 2019; Lorenz et al., 2017; Ohira & Suzuki, 2024; Sas-Chen et al., 2020; Shigi, 2018; Tsai et al., 2025). The chemical stability and structural plasticity conferred by tRNA modifications likely play roles in enabling translation to proceed under these particularly extreme physicochemical conditions.

High-throughput tRNA sequencing has become a powerful tool for quantifying tRNA abundance and expression dynamics, complementing data from traditional mRNA and small RNA-seq approaches that were not designed to measure highly structured and modified RNAs reliably. In addition to profiling tRNA expression, specialized tRNA-seq methods allow the detection of chemical modifications by leveraging the tendency of certain base modifications to disrupt Watson-Crick (W-C) pairing during reverse transcription (RT), which can be captured by quantifying the precise positions of misincorporations or early termination of resulting cDNA strands (Behrens et al., 2021; Cozen et al., 2015; Erber et al., 2020; Gustafsson et al., 2022; Kimura et al., 2020; Nakano et al., 2025; Upton et al., 2021; Zheng et al., 2015). While mass spectrometry remains the gold standard for the direct chemical identification of modified nucleosides, RT-based sequencing offers a complementary, high-throughput, low-cost, nucleotide-resolution approach ideal for large-scale comparative studies.

One of the newest RT-based RNA-seq methods applied to tRNAs, Ordered Two-Template Relay sequencing (OTTR-seq), enhances detection of modification-sensitive misincorporation events while maintaining full-length tRNA coverage (Upton et al., 2021; K. Zhang et al., 2025). OTTR-seq utilizes a processive reverse transcriptase with a defined template-switching mechanism, thereby enhancing the uniformity of read coverage across structured RNAs. This makes OTTR-seq particularly well-suited for transcriptome-wide profiling of tRNA modifications, where detecting both conserved and context-dependent modification events requires consistent capture of intact, full-length tRNAs. In archaeal species, where tRNAs are highly structured and often densely modified, this approach enables a comprehensive view of the modification landscape, revealing relationships between modification frequency, tRNA sequence, and enzyme specificity.

Beyond cataloging modifications, understanding the evolution of tRNA-modifying enzymes allows for insights into the rules governing substrate selection. These enzymes can share conserved structural motifs but often exhibit variable domain architectures and phylogenetic distributions, complicating predictions of substrate specificity (Hori, 2023; Jackman & Alfonzo, 2013). Few enzyme-target relationships are governed not only by conserved nucleotide motifs (e.g., G10oU25 pairs) but also by lineage-specific structural features and anti-determinants that influence accessibility or folding of target regions (Helm, 2006; Motorin & Helm, 2011).

Much like each aminoacyl-tRNA synthetase (aaRS) recognizes and charges particular subsets of tRNAs based on distinct sequence and structural identity elements (Westhof et al., 2022), tRNA modification enzymes recognize a combination of specific sequence motifs and/or structural elements shared by their target tRNAs. Some modifications, such as N2-methylguanosine (m^2^G) or N2, N2-dimethylguansine (m^2^ G) at tRNA nucleotide position 10 at the base of the D-stem, are essential early in pre-tRNA maturation, mediating the correct folding of the L-shape tertiary structure (Hori, 2019; Shigi et al., 2006). Transfer RNA modification pathways are interconnected and exhibit cooperativity with one another, which, in studied cases, influences stress response or leads to disease through modulation of tRNA stability and decoding behavior (Hernandez-Alias et al., 2022; Sokołowski et al., 2018). To add to this complexity, the precise target specificity of tRNA modification enzymes evolves across diverse phyla, causing uncertainty or inconsistencies between studies on the same family of modification enzymes in different model systems (de Crécy-Lagard et al., 2019).

Among the various tRNA modification types, methylation is the most prevalent, with at least 13 distinct chemical variations of methylated RNA bases shared among all three domains of life (Hori, 2014; Jackman & Alfonzo, 2013; Motorin & Helm, 2011). Despite this overlap, modification enzymes vary widely in their protein domain architectures and substrate specificities across the tree of life. While some, like TrmI, function as multi-subunit complexes in eukaryotes but as homotetramers in Archaea (Roovers et al., 2021), others show variability in domain architectures that influence tRNA recognition (Dixit et al., 2019). These differences impact how modification enzymes selectively recognize and act on diverse tRNA sequences, yet the extent of this variation and its functional consequences remain poorly understood.

Here, we present a comprehensive analysis of archaeal tRNA modifications, revealing novel clade-specific patterns shaped by environmental adaptation and mirrored in enzyme divergence. Using OTTR-seq across nine species, including thermophiles, methanogens, acidophiles, and halophiles, we chart the landscape of RT-detectable tRNA modifications to investigate how modification dynamics evolve across distinct ecological niches. To correlate these new patterns to their enzymes, we examined four AdoMet (SAM)-dependent methyltransferases that target key positions: Trm14 (m^2^G/m^2^ G at position 6), Trm11 (m^2^G/m^2^ G at position 10), Trm1 (m^2^G/m^2^ G/m^2^ Gm at position 26), and Trm10 (m^1^A9/m^1^G9). These enzymes span two major classes, with Trm14, Trm11, and Trm1 belonging to class I Rossmann-fold MTases and Trm10 to the class IV SPOUT family (Bujnicki et al., 2002). Our integrated analyses highlight position-specific and isoacceptor-dependent variations, coupled with changes corresponding to presence/absence of modification enzyme orthologs or changes to their domain architectures. Together, these data uncover strong co-variation between enzyme families and modifications at key tRNA positions, underscoring the deep co-evolution of archaeal tRNAs with their modifying enzymes, and bootstrapping a probabilistic framework for prediction of modification presence or absence directly from genome sequence.

## Results

### Detection of base-specific misincorporations reveals tRNA modification dynamics across archaea

We profiled post-transcriptional tRNA modifications across nine archaeal species (Table 1) using OTTR-seq (Upton et al., 2021) to identify reverse transcription (RT)-induced misincorporation signatures of tRNA modifications (Figure 1). Known modification sites in six of these species (*H. volcanii, M. jannaschii*, *M. maripaludis*, *P. furiosus, S. acidocaldarius,* and *T. kodakarensis*) (Gupta, 1984; Hirata et al., 2019; Wolff et al., 2023; Yu et al., 2019) provided a validated reference for evaluating modification detection (Additional file: Table S0, S1), while additional comparisons with *Thermococcus sp.* AM4 and *Sulfolobus islandicus* enabled assessment of short-term evolutionary conservation (Figure 1A). We predicted sites of tRNA modification via prominent and statistically significant misincorporation (MI) frequencies, defined as the proportion of sequencing reads at a given nucleotide position that differ from the reference base. Misincorporation frequencies define the extent to which a modification disrupts the incorporation of the cognate nucleotide during reverse transcription, typically by disrupting Watson-Crick (W-C) base pairing, and provide a general proxy for the presence and stoichiometry of modification events. To evaluate detection confidence, we established two key metrics that combine statistical significance and MI frequency thresholds to control both the sensitivity and specificity of modification detection (see Methods). Using this system, the distribution of high- and moderate-confidence modifications is summarized by position (Figure 1B) and across all tRNAs (Additional file: Fig. S1 and Fig. S2, Table S1).

**Table 1.** Summary of study species and relevant attributes.

| Species (abbr) | Phylum | Class | Habitat | Temperature / pH Preference | Number of tRNA Genes |
| --- | --- | --- | --- | --- | --- |
| *Sulfolobus islandicus* (Sis) | Crenarchaeota | Thermoprotei | Volcanic hot springs | Thermoacidophile (75°C, pH 2-3) | 45 |
| *Sulfolobus acidocaldarius* (Sac) | Crenarchaeota | Thermoprotei | Terrestrial hot springs | Thermoacidophile (75°C, pH 2-3) | 46 |
| *Pyrococcus furiosus* (Pfu) | Euryarchaeota | Thermococci | Hydrothermal marine sediments | Hyperthermophile (95°C, pH 6-7) | 46 |
| *Thermococcus* sp. AM4 (AM4) | Euryarchaeota | Thermococci | Sea hydrothermal vents | Hyperthermophile (80°C, pH 6-7) | 46 |
| *Thermococcus kodakarensis* (Tko) | Euryarchaeota | Thermococci | Sea hydrothermal vents | Hyperthermophile (85°C, pH 6-7) | 46 |
| *Halobacterium salinarum* (Hsa) | Euryarchaeota | Halobacteria | Hypersaline lakes and salterns | Extreme halophile (40°C, pH 7) | 47 |
| *Haloferax volcanii* (Hvo) | Euryarchaeota | Halobacteria | Hypersaline environments | Moderate halophile (42°C, pH 7) | 47 |
| *Methanococcus maripaludis* (Mma) | Euryarchaeota | Methanococci | Salt marshes | Mesophile (37°C, pH 7-8) | 35 |
| *Methanocaldococcus jannaschii* (Mja) | Euryarchaeota | Methanococci | Marine hydrothermal vent | Hyperthermophile (85°C, pH 6-7) | 34 |

**Figure 1.**
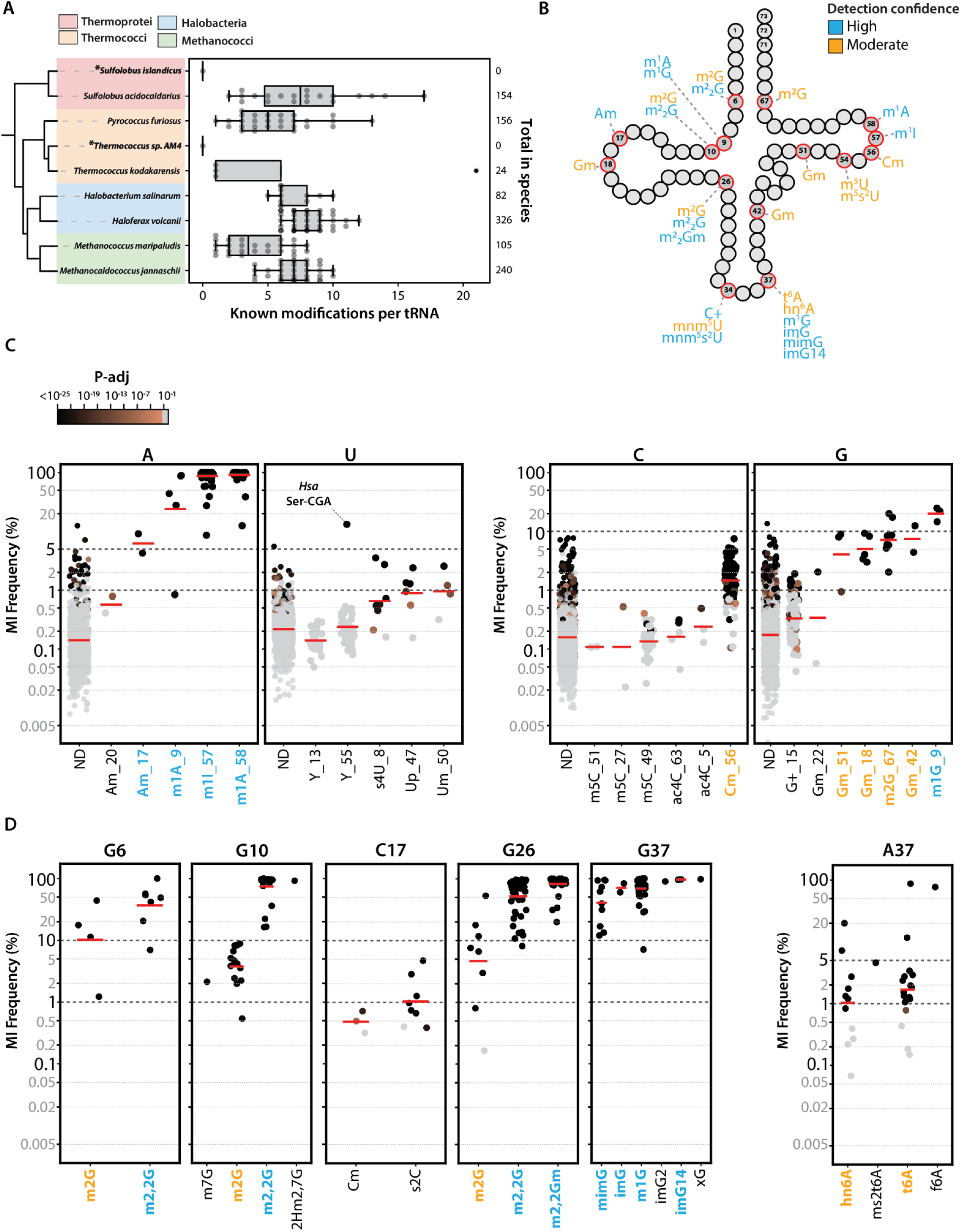
Known archaeal tRNA modifications and their misincorporations (MI) rates. A) Phylogenetic tree of representative archaeal species, annotated in boxplots, where each datapoint represents a given tRNA with annotated modifications. The number of known tRNA modifications per species was curated from Modomics and literature sources. Species wherein no literature sources define tRNA modifications are marked with an asterisk(*) B) Cloverleaf schematic of tRNA secondary structure indicating detection confidence level at positions of known tRNA modifications across all species. Modifications are color-coded by estimated confidence of detection via OTTR-seq, based on average misincorporation frequency (MI) and adjusted p-values. High confidence detection (blue) was classified by base-specific MI rates (mean MI > 10% for cytosine and guanine; mean MI >5% for adenine and uracil) and an adjusted p-value <0.05. Modifications with significant p-values but misincorporation rates below the respective base threshold were classified as moderate-confidence (orange). C) Log-scale position-specific MI frequencies stratified by nucleotide identity for known modifications and a set of sites classified as ND (Not Determined). ND denotes positions where there is currently no published data supporting a particular modification in the specific species and tRNA context considered (see methods). Each point represents a position in a specific tRNA, and red lines show mean MI values. Adjusted p-values indicate statistical support for misincorporation (copper to black gradient indicates p-adj < 0.05; p-adj > 0.05 in grey). D) MI profiles at selected positions where multiple known modification types can occur from the same base (e.g. G6, G10, C17, G26, G37, A37). Each subpanel shows MI distributions stratified by specific chemical modification. Differences in MI frequencies aid in distinguishing closely related modification types such as m^2^G vs. m^2^ G at G10. Confidence in detection varies across positions and modification types.

These MI-detectable tRNA modifications exhibited a range of misincorporation profiles, as categorized by nucleotide (Figure 1C). We note that relative to adenine and uridine, guanine and cytosine exhibit higher background misincorporation rates for unmodified tRNA positions, which required adjusted MI thresholds for our detection confidence classification (see Methods). Modifications disrupting the W-C face (e.g., m^1^A, m^1^I, m^2^G, m^2^ G) consistently showed the highest MI rates, and these deviations involving position-specific and/or isotype-specific patterns revealed biologically relevant differences. For example, while m^1^A58 typically results in near-complete misincorporation, Sulfolobales species showed reduced MI rates at A58 in tRNA^Ala^ (Additional file: Fig. S4). Despite lower MI rates, the read identity (patterns of misincorporation) and read depth supported m^1^A modification at this position, suggesting substoichiometric modifications and apparent tRNA-specific regulation (Figure 1C, Additional file: Fig. S3). Similar observations were made for A57 in *P. furiosus* Thr^GGU^ where rates of misincorporation markedly dropped, but the patterns of misincorporation were consistent with m^1^I (Additional file: Fig. S4).

Other notable outliers included pseudouridine at position 55, which was consistently undetectable across species, as expected given its minimal impact on reverse transcription. However, in *H. salinarum* Ser^CGA^, we observed an unusually high misincorporation rate and a significant p-value for modification at this position. While pseudouridine alone is unlikely to account for this signal, it may reflect the presence of a modified derivative such as m^3^U or m^3^Ψ, which can disrupt the W-C face but has not previously been identified at position 55 (Additional file: Fig. S4).

An important observation, consistent with prior tRNA-seq studies (Behrens et al., 2021; Nakano et al., 2025; K. Zhang et al., 2025), was that different modification types occurring at the same position and base had characteristic misincorporation behaviors (Figure 1D, Additional file: Fig. S1). The relative distributions of these rates varied by modification type from the same position. In most cases, this allowed us to make reasonable predictions, choosing between known modifications at the same position (Additional file: Table S1). For example, at position 10, the misincorporation frequency for m^2^ G was consistently higher than that for m^2^G, allowing for distinction between these modifications. When these patterns are augmented by known positives in previously analyzed species (Figure 1C), as discussed in the next section, this becomes a powerful method for screening many tRNAs for all modifications that cause misincorporations.

### Homology-based predictions of tRNA modifications identify both conserved and dynamic sites of modification

Mapping known modification sites to positions of misincorporations in previously characterized species allowed us to predict their presence in related species lacking experimental annotations (Figure 2). For example, the known modification profile of tRNA^iMet^ in *P. furiosus* was mapped to concordant sites of misincorporation in *T. kodakarensis*. In this case, we were able to predict all known and detectable modification sites in *T. kodakarensis*, except for 2’-*O*-methylguanine (Gm) at position 22 (Figure 2A), which has a relatively low misincorporation frequency, even in the reference *P. furiosus* tRNA (Figure 1C). Additional pairwise comparisons across species further supported the predictive power of misincorporation-based inference of site-specific tRNA modifications. Across comparisons, known and predicted misincorporation frequencies of modified homologous tRNAs were highly correlated (Additional file: Table S3; see Methods). For species within the same Class (e.g., Thermococci), Pearson correlation coefficients ranged from 0.87 to 0.95 (Figure 2B-F, Additional file: Fig S5), and this strong agreement indicates that modified homologous tRNAs frequently share conserved misincorporation patterns, reinforcing the value of using known sites of tRNA modification as predictors for tRNA modification patterns in closely related taxa.

**Figure 2.**
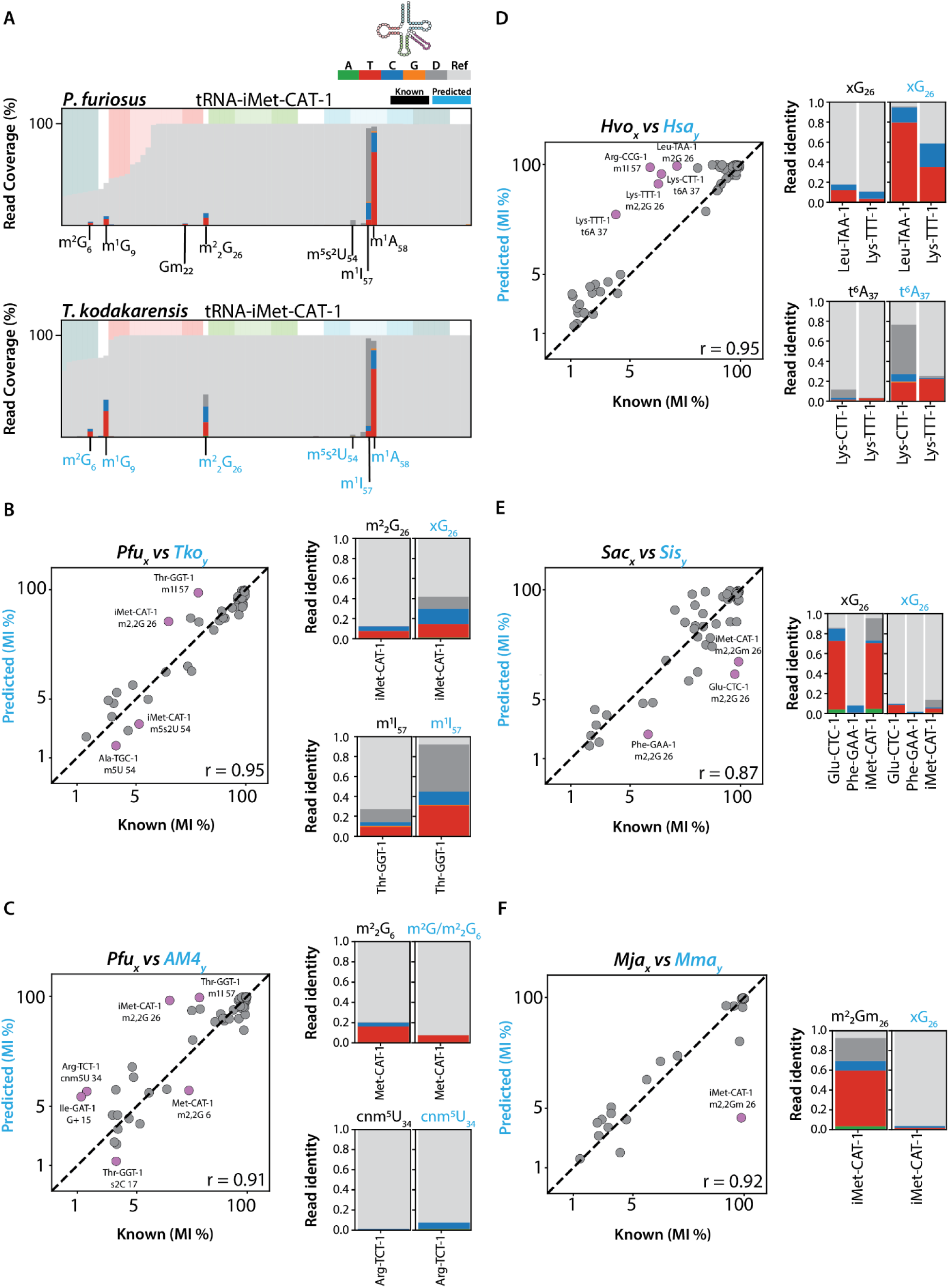
Overview of homology-based tRNA modifications predictions. A) Representative example of homology-based predictions for the initiator methionine (tRNA^iMet^) in *P. furiosus* (top) and *T. kodakarensis* (bottom). The coverage plot depicts the known modification sites indicated (e.g. black) and newly predicted sites highlighted (blue). Mapped-read coverage plot illustrates the read depth and pinpoint sites of modification. B-F) Pearson correlation between known modifications vs homology-predicted modifications. Each point in the scatter plot corresponds to a specific modification type and site from homologous tRNAs. The diagonal line represents perfect correlation (r=1) between known and predicted MI. Outliers (± 2 SD) are highlighted (purple) and annotated with their corresponding tRNA, modification, and position. The read identity (pattern of misincorporation) for select outliers are depicted next to the scatter plot, with known modification position (black) compared to newly predicted (blue) (See all outliers in Supplemental Figures) B) *P. furiosus (Pfu)* vs *T. kodakarensis (Tko)*. Select outliers iMet^CAU^ m^2^ G at position 26 and Thr^GGU^ m^1^I at position 57. C) *P. furiosus (Pfu)* vs *T. species* AM4 *(AM4).* Select outliers Met^CAU^ m^2^ G at position 6, and Arg^UCU^ cnm^5^U at position 34. D) *H. volcanii (Hvo)* vs *H. salinarum (Hsa)*. Select outliers Leu^UAA^ m^2^G, Lys^UUU^ m^2^ G at position 26, and Lys^CUU^, Lys^UUU^ t^6^A at position 37. E) *S. acidocaldarius (Sac) vs S. islandicus (Sis).* All outliers Glu^CUC^ and Phe^GAA^ m^2^ G, and iMet^CAU^ m^2^ Gm at position 26. F) *M. jannaschi (Mja)* vs *M. maripaludis (Mma).* Outlier iMet^CAU^ m^2^ Gm at position 26. For positions with more than one modification type (e.g., m^2^G/m^2^ G) “x” was used. This notation is also used for most predictions, as exact modification types weren’t ascertained.

However, deviations from expected misincorporation patterns revealed additional layers of biological variation (Figure 2B-F, purple off-diagonal points). These unexpected profiles were particularly informative, suggesting partial or context-specific modifications in homologous tRNAs between species within the same Class, family, or genus. Notably, for many published modification data sets, there is often no discrimination between fully and partially modified nucleotides, leading to ambiguity when comparing them to our MI frequencies. For example, the known m^2^ G in iMet^CAU^ of *P. furiosus* had a lower and more variable pattern of misincorporation compared to the predicted modification of the homologous tRNA in *T. kodakarensis* at position 26 (Figure 2B; Additional file: Fig. S6). Interestingly, the expected modification in *T. kodakarensis* had a larger proportion of RT-induced deletions (Figure 2B, m^2^ G v *x*G). While the triple methylation of guanine (m^2^ Gm) is often associated with deletions and higher rates of misincorporation, we broadly define all likely G-derived modifications at position 26 as “*x*G”, as there are usually two or more modification types that occur at this position, which cannot be consistently distinguished solely through MI frequencies. Similarly, the difference observed at position 57 in Thr^GGU^ between these same two species shows that the patterns of misincorporation are consistent with m^1^I despite variation in rates of misincorporation.

This pattern of divergence was also observed in comparisons with other species. In *P. furiosus* vs *T. sp.* AM4, we identified additional outliers (Figure 2C, purple points), including the m^2^ G in iMet^CAU^ at position 6, a site known to have either m^2^G or m^2^ G in other tRNAs. The relatively low misincorporation frequency in *T. sp.* AM4 aligns more closely with m^2^G, prompting us to conservatively annotate this site as m^2^G/m^2^ G. Another notable outlier in this comparison was position 34 in Arg^UCU^, where the misincorporation pattern in *T. sp.* AM4 was consistent with cnm^5^U, despite being near the lower limit of detection (∼1% MI). This highlights the potential to identify U-derived modifications at the wobble position, even when present at low levels, though such predictions warrant cautious interpretation.

Further comparisons between *H. volcanii* and *H. salinarum* (Figure 2D) revealed similar variations in modification state, which resulted in unresolved prediction ambiguity. In this case the differences in stoichiometry suggest that the “known” reference site may merit re-examination. In *H. volcanii*, the low MI rate and read-identity pattern in Lys^UUU^ are more consistent with m^2^G, matching the known m^2^G in Leu^UAA^. This does not preclude a minority m^2^ G fraction in Lys^UUU^ that may explain the reference (Gupta, 1984). By contrast, in *H. salinarum* both Leu^UAA^ and Lys^UUU^ exhibit the elevated MI typical of m^2^ G. Because the reference datasets do not report stoichiometry or mixed populations, we conservatively annotate these calls. Additionally, modifications at position 37, such as N^6^-threonylcarbamoyladenosine (t^6^A) in Lys^CUU^ and Lys^UUU^, showed variations in predictions, reflecting either partial modification or differences in species-specific regulation.

A similar trend emerged in *S. acidocaldarius* versus *S. islandicus* analyses (Figure 2E).

Modifications at position 26 in Glu^CUC^, Phe^GAA^, and iMet^CAU^ were all underrepresented in *S. islandicus,* though read identity patterns remained consistent with the expected modification types. These findings suggest that the stoichiometry of modified versus unmodified, rather than the type of modification, may vary between these species. Finally, in *M. jannaschii* vs *M. maripaludis* (Figure 2F), position 26 in iMet^CAU^ was again identified as an outlier. In this case, the hyperthermophile (*M. jannaschii*) has MI patterns consistent with m^2^ Gm, whereas the mesophile (*M. maripaludis*) has patterns consistent with m^2^G.

Together, conserved modifications at positions like 6, 26, and 37 appear to be hotspots for interspecies variation in stoichiometry or type. These comparisons highlight how deviations from expected misincorporation frequencies can predict species-specific differences in modification state. Moreover, read identity patterns support the biological plausibility of these differences, as distinct patterns of misincorporation are often consistent with those of known modifications. Our observations emphasize the caution required in homology-based modification predictions, where careful contextual interpretation is required. For our downstream analyses, we focus on modifications that have moderate to high confidence of detection, and use xN notation at positions where the exact modification type is unable to be inferred.

### Clade-specific patterns reveal coordinated modification in the acceptor stem and core regions

Distinct modification patterns in the acceptor stem and tRNA core regions reveal coordinated signatures that vary by phylogenetic clade. In particular, m^2^G and m^2^ G modifications at positions 6 and 67 (base pairing in the acceptor stem) are frequently predicted in hyperthermophilic species (Figure 3A). While prior studies have reported individual modifications at these positions in select tRNAs from *M. jannaschii* and *P. furiosus* (Wolff et al., 2023; Yu et al., 2019), the broader pattern of coordination has not been previously recognized.

**Figure 3.**
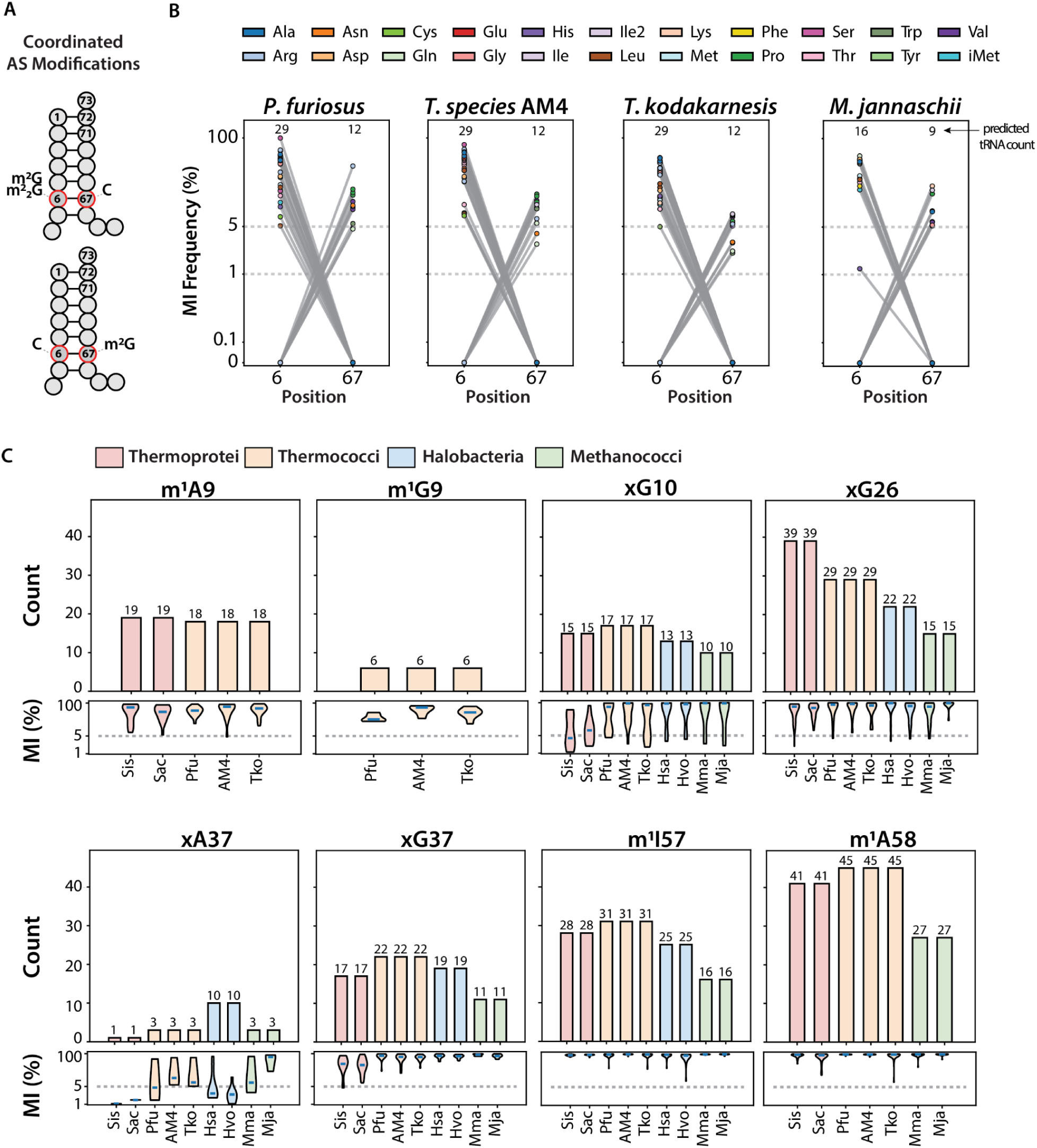
Clade-specific coordination patterns in the acceptor stem and conserved clade-specific tRNA modifications. A) Schematic representation of coordinated modification patterns at the acceptor stem (AS) of the tRNA cloverleaf secondary structure, highlighting a mutually exclusive relationship between position 6 (typically m^2^G or m^2^_2_G) and position 67 (m^2^G). Observed across multiple hyperthermophiles, the data suggest that these modifications do not co-occur within the same tRNA, possibly due to structural constraints. B) Paired MI frequencies for position 6 and 67 across four hyperthermophilic archaea: *P. furiosus, T. sp.* AM4*, T. kodakarensis,* and *M. jannaschii.* Each line connects paired MI values for individual tRNAs, with colored dots indicating isotype identity. Only moderate and high-confidence predictions (adjusted p-value < 0.05) are shown. Numbers above each panel indicates the total number of predicted modified tRNAs at positions 6 and 67, respectively. A pattern of coordinated modification is consistently observed. C) Distribution of MI frequencies for select moderate to high-confidence tRNA modifications shared across archaeal clades, grouped by phylogenetic class: Thermoprotei (salmon), Thermococci (tan), Halobacteria (blue), and Methanococci (green). Each violin plot summarizes the frequency and variation of a specific modification (e.g. m^1^A9, m^1^G9, m^2^G10, etc.) across species, with counts (above plots) indicating the number of modified tRNAs per group. This highlights the phylogenetic conservation and variability of tRNA modification patterns across lineages.

Our expanded analysis reveals that across multiple species, position 6 is more frequently modified than position 67, with misincorporation rates suggesting a predominance of m^2^_2_G6 over m^2^G6. In contrast, m^2^G67 is less frequent but consistently observed in distinct tRNA subsets (Additional file: Fig. S2 and Table S4). The two modifications are not predicted to co-occur within the same tRNA (Figure 3B), indicating a likely mutually exclusive pattern of modification. This trend is particularly evident in all Thermococci species and *M. jannaschii* (Methanococci). Most G6 conserved tRNAs are predicted to have a modification across these species (Additional file: Fig. S2 and Table S4). However, certain tRNAs in Thermococci, such as Glu^UUG^ and Ile^GAU^, possess a C6 (and m^2^G67), precluding the possibility of m^2^G or m^2^ G at this position.

Coordination of modification patterns is also observed between positions 10 and 26, two highly conserved sites of modification across archaeal tRNAs. While the predicted modifications conserved in species from Halobacteria and Methanococci tend to occur in mutually exclusive subsets of tRNAs, co-occurrence of modifications at both sites within the same tRNA is found in Thermococci and Sulfolobales (Additional file: Fig. S7 and Table S5). This pattern is especially pronounced in Sulfolobales, where multiple tRNAs such as tRNA^Asp^, tRNA^Glu^, and tRNA^Val^ are predicted to carry both m^2^G/m^2^ G10 and m^2^ G26 (Additional file: Fig. S2, S7 and Table S5). In contrast, while Thermococci also exhibits co-modification of position 10 and 26, their modification types differ. With lower rates of misincorporation at position 10, their tRNAs may instead carry m^2^G10 and m^2^ G26 (Additional file: Fig. S7). These differences are not entirely surprising as these specific clades span two phyla (Crenarchaeota and Euryarchaeota). Their preferences in patterns of modifications suggest evolutionary divergence in enzymatic activity and substrate selection.

Building on these observations, coordinated modification patterns at 10 and 26 often correspond to specific sequence contexts within tRNAs. Predicted G10 substrates typically base pair with U25, consistent with previous observations (Urbonavičius et al., 2006), while G26 modifications tend to occur in tRNAs lacking this pairing (Additional file: Fig. S8, Table S7). Interestingly, in Sulfolobales, several tRNAs predicted to be modified at both positions contain G10oU25 but lack the D-arm motifs (e.g., U13oU22, U13-A22) commonly associated with m^2^G10/m^2^ G10 in other clades. These findings suggest that the coordination of modifications at structurally proximal positions is reflecting shared structural constraints across archaeal tRNAs, while clade-specific tuning of enzyme-substrate interactions contributes to lineage-specific nucleotide variation.

### Conserved and clade-specific tRNA modifications at functional positions support distinct evolutionary adaptations

Several tRNA positions display broad conservation across archaeal species, although their distribution and frequency vary by lineage and environmental niche (Figure 3C, Additional File: Fig. S2). Modifications at position 10 are widespread across clades, reflecting their proposed functional role in reinforcing the D-stem (Urbonavičius et al., 2006). Our predictions show the frequency of *x*G10 (m^2^G or m^2^ G) is slightly more prevalent in both Thermococci and Sulfolobales, where over a third of the conserved orthologous tRNAs are modified (Figure 3C, Additional File: Fig. S2). While previous work has shown the importance of m^2^ G10 for species growing at elevated temperatures (Gabant et al., 2006), we find that it is also a common modification in mesophilic Halobacteria (28% of tRNAs in *H. volcanii*) and Methanococci (27% of tRNAs in *M. maripaludis*). Interestingly, in Thermococci and Sulfolobales, several tRNAs such as Gly^UCC^ and tRNA^Val^ are predicted to carry *x*G10 despite lacking canonical sequence features generally associated with this modification, such as the G10oU25 pair (Additional File: Table S6).

Position 26 is one of the most consistently modified sites across archaeal species, with nearly all clades displaying high numbers of modified tRNAs (Figure 3C). Sulfolobales show the highest counts with nearly all tRNAs predicted with *x*G26 (m^2^G, m^2^ G, m^2^ Gm), followed by Thermococci, Halobacteria, and Methanococci. While previous work has linked the prevalence of m^2^ G26 and its hypermodified derivative m^2^ Gm with thermophiles (Kowalak et al., 1994; Wolff et al., 2023), our results show highest modification frequency in acidophiles, indicating that *x*G26 serves functions beyond thermal adaptation alone. Importantly, not all G26-containing tRNA are modified, particularly in Halobacteria, where iMet^CAU^ and Gly^GCC^ are not predicted to be modified, implying selective targeting by Trm1. In several lineages, predicted *x*G26 modifications appear mutually exclusive with G10oU25 base pairs, consistent with previously defined anti-determinants (Urbonavičius et al., 2006), though we find that Sulfolobales frequently bypass this rule (e.g., tRNA^Glu^ and tRNA^Phe^; Additional File: Fig. S8).

Our results reveal additional patterns that suggest clade-specific evolutionary pressures beyond thermophily alone. For instance, position 9 shows clear phylogenetic divergence (Figure 3C). Sulfolobales predominantly carry m^1^A9 across nearly all A9-containing tRNAs, whereas Thermococci harbors both m^1^A9 and m^1^G9 (Additional File: Fig. S2), reflecting broader substrate recognition by their Trm10 homologs (Krishnamohan & Jackman, 2019). Notably, neither Halobacteria nor Methanococci exhibit evidence of position 9 modifications, consistent with the absence of detectable Trm10 homologs in these clades.

Modifications at position 37, a universally functional site adjacent to the anticodon, display substantial chemical diversity across archaea. Predicted *x*A37 modifications (typically t^6^A and its derivatives) are largely restricted to Halobacteria, however these are often called with low confidence in our experiments and may not reflect true variations between clades (Additional File: Fig. S1 and Fig. S2). In contrast, *x*G37 modifications (m^1^G and wyosine-like derivatives) are confidently predicted and observed to be most prevalent in Thermococci, with nearly half of the conserved orthologous tRNAs being modified. Interestingly, Gly^UCC^ in *P. furiosus* uniquely has a modified G37 (all other species contain A37) and tRNA^Val^ displays frequent G37 and A37 variations from species to species. The distinct distributions of these support clade and species-specific strategies for tuning decoding accuracy and anticodon loop stability (de Crecy-Lagard et al., 2010; W. Zhang & Westhof, 2025).

Conserved T-arm modifications at position 57 and 58 are among the most consistently observed modifications across species (Figure 3C, Additional File: Fig. S2). Both m^1^I57 and m^1^A58 are most frequent in Thermococci and Sulfolobales, supporting their proposed role in stabilizing tRNA tertiary structure under extreme conditions (Roovers et al., 2021). In most lineages, m^1^A58 occurs in all A58 tRNAs, with the exception of Halobacteria which lack this near-universal modification. While m^1^I57 is also widespread, occurring in all A57 tRNAs, its occurrence varies from species to species and even within isoacceptors. For instance, Val^UAC^ in *P. furiosus* has G57, consistent with all tRNA^Val^ among Thermococci, but distinctly has A57 in Val^CAC^ and Val^GAC^ which is modified with m^1^I.

Together, these observations outline a backbone of the archaeal tRNA modification landscape, highlighting core functional sites discernable with our high-throughput method (positions 9, 10, 26, 37, 57, and 58). Patterns of variation may help accommodate structural changes needed due to environmental pressures, ranging from temperature and pH to salinity. The balance between conservation and diversification may help to sustain key modification roles while also allowing enough flexibility for unique and specialized functional roles to undergo selective adaptation. From an RNA modification-based regulation perspective, these sites are of particular interest because they demonstrate key design axes for biologically or synthetically altered tRNA function. Tracing these altered patterns of modification back to the source enzymes becomes a feasible goal with the scale of data gathered in this study.

### Phylogenetic variation in tRNA methyltransferase architectures reveals enzyme specialization across archaea

We carried out comparative analysis of tRNA methyltransferase ortholog families to examine enzyme distribution and domain architecture variation alongside clade-specific modification patterns (Figure 4). We focused on four AdoMet-dependent MTases known to catalyze methylation at specific tRNA positions that showed clade-specific patterns examined above.

**Figure 4.**
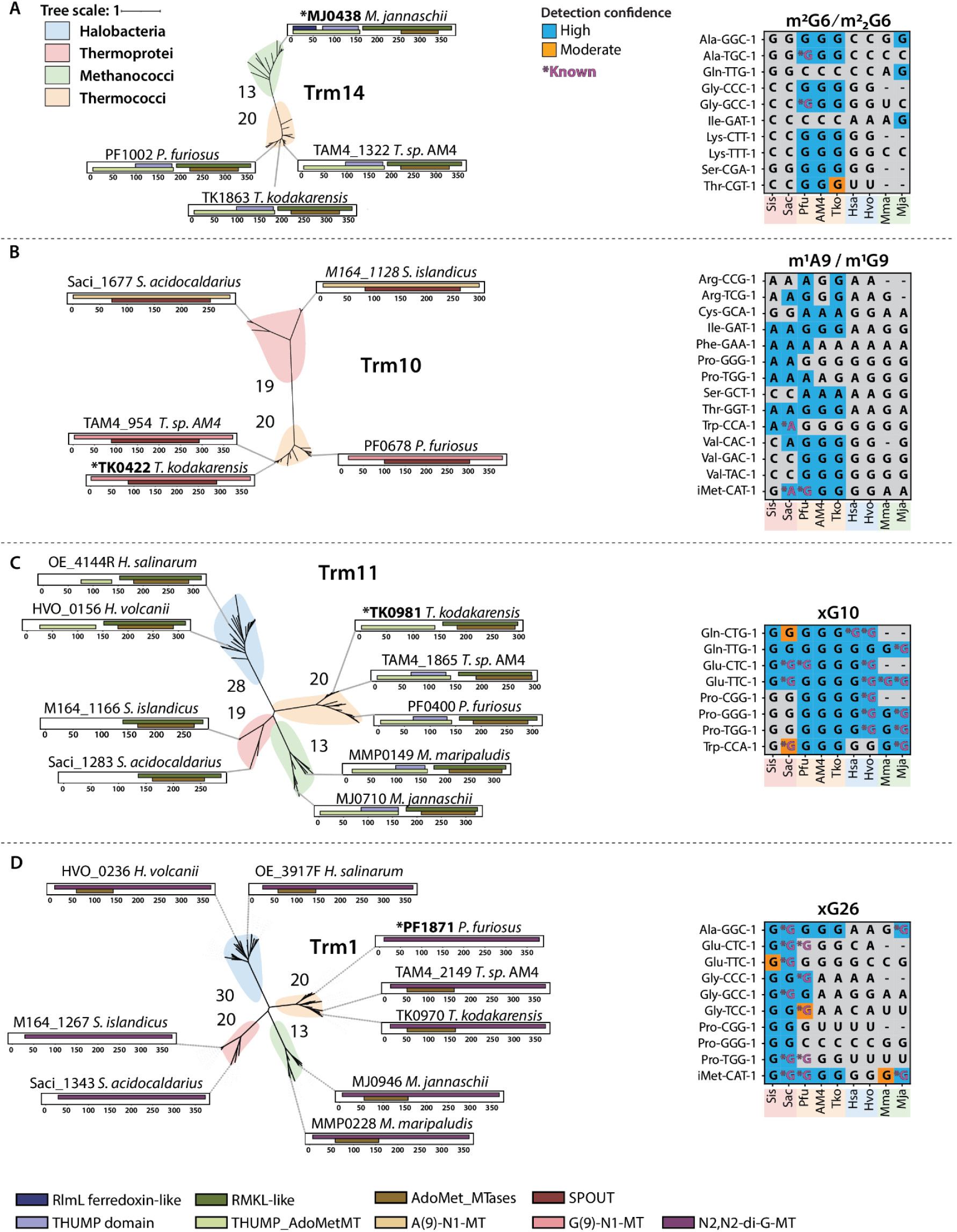
Phylogenetic and structural diversity of predicted tRNA modification enzymes and their target substrates across Archaea. Each panel presents a phylogenetic tree of predicted homologs of a specific tRNA-modifying enzyme (left), alongside a heatmap summarizing the confidence level of predicted substrates across species (right). Trees are constructed from clustered protein sequences colored by archaeal clades: Halobacteria (blue), Thermoprotei (salmon), Methanococci (green), and Thermococci (tan). Domain architecture annotations are shown on the tree branches, indicating key functional domains for three class I MTases (Trm14, Trm11, Trm1) and a class IV SPOUT MTase (Trm10). A) Trm14 homologs are restricted to Methanococci and Thermococci and are associated with predicted m^2^G6/m^2^ G6 modifications in specific tRNA substrates. The corresponding heatmap shows predicted modifications for representative tRNAs across genomes. B) Trm10 homologs are exclusive to Thermoprotei and Thermococci and are associated with m^1^A9 or m^1^G9 modifications. The heatmap displays inferred substrate preferences for these homologs across tRNAs and illustrates the dual-functional properties unique to Trm10 in Thermococci. C) Trm11 homologs are broadly conserved, although THUMP domain truncations are observed in Thermoprotei. These homologs are associated with m^2^ G10 modifications in select substrates. Divergence is most notable in tRNA^Pro^. D) Trm1 homologs show widespread conservation, with Thermoprotei variants displaying divergence in the AdoMet domain. Predicted modification patterns at position 26 include m^2^ G and other variants such as m^2^G/m^2^ Gm (*x*G26). Positions previously known and annotated as m^2^G reflect sites with adjusted p-values > 0.05, indicating lower statistical support. Together, these patterns highlight coordinated evolution of enzyme domain architectures and their target specificity across archaeal lineages.

Trm14 (m^2^G6/m^2^ G6), a THUMP (Thiouridine synthases, RNA methylases and Pseudouridine synthases)-related methyltransferase, occurs exclusively in hyperthermophiles and retains the canonical domains consistent with previously characterized structures (Fislage et al., 2012; Menezes et al., 2011) (Figure 4A, Additional File: Table S8). Subtle N-terminal differences distinguish clades (e.g., MJ0438 in *M. jannaschii* uniquely has a hit for an RlmL-like domain), however, variations in target substrate are better explained by distinct tRNA features between these species (Figure 4A). While Trm14 is known to target G6, methylation of G67 remains undetermined. There is evidence of Trm14 involvement in m^2^G67 (Hirata et al., 2019), yet a survey of paralogs identified candidates within these clades that may also be involved (Additional File: Table S12).

Contrary to earlier impressions of broad archaeal conservation (Krishnamohan & Jackman, 2019), Trm10 orthologs appear narrowly conserved in Thermococci and Thermoprotei (Figure 4B, Additional File: Table S12). Members of the SPOUT-family are notoriously challenging to model based on primary sequence, however there are detected features compatible with bifunctional G9/A9 N1-methylation in Thermococci, consistent with *T. kodakarensis* TK0442’s dual specificity (Krishnamohan et al., 2019) (Figure 4B, Additional File: Table S9). In Sulfolobales, predicted domain features diverge toward A9-specific residues, and the misincorporation profiles shift accordingly toward A9 targets (Figure 4B), indicating a phylogenetic split between bifunctional and A-specific Trm10s.

Trm11 (m^2^G10/m^2^ G10) is broadly distributed, but Sulfolobales diverge from the canonical domain organization described in *T. kodakarensis* TK0981 (Hirata et al., 2016; Urbonavičius et al., 2006). While they retain the C-terminal catalytic domain, they lack the expected N-terminal THUMP tRNA-binding domain (Figure 4C, Additional File: Table S10). This architectural loss coincides with a shift in substrate that co-varies with specific isoacceptors (notably tRNA^Pro^) in our predictions (Figure 4C). Given there are many known partnership with Trm112 in *H. volcanii* (van Tran et al., 2018), a modular binding partner (Trm11-Trm112 or analogous factor) may compensates for the missing THUMP in Sulfolobales.

Trm1 (m^2^G26/m^2^ G26) lacks a THUMP domain (Hori, 2023; Ihsanawati et al., 2008) yet, like Trm11, catalyzes mono-then dimethylation. Canonical domain organization is broadly conserved for all predicted orthologs, although profile matches for SAM motifs in the N-terminal region diverge in Sulfolobales (Figure 4D, Additional File: Table S11). Despite subtle differences in architecture, target usage diverges significantly in Sulfolobales, where anti-determinant rules are broken. For instance, both *x*G10 and *x*G26 are predicted to co-occur in tRNA^Glu^ (Figure 4C and 4D).

Together, our comparison of these key enzyme families establishes a predictive link between domain architecture and target nucleotide choice, further supporting a functional split in Trm10 specificity, and revealing a new THUMP-independent Trm11 with isoacceptor bias. This comparative methodology can be applied to other tRNA modification enzymes extending beyond methyltransferases (e.g., pseudouridine synthases), as more data is available and methods in high-throughput modification detection continue to improve.

### tRNA sequence and structure features reveal determinants and anti-determinants of modification enzyme targeting

Positions with predicted modifications correspond to conserved nucleotide features in specific tRNAs across various species. Likewise, shifts in predicted modifications are linked to changes in nucleotides at distinct positions. Our comprehensive analysis across multiple archaeal clades expands upon previous observations, reinforcing and broadening the understanding of tRNA features critical for modification enzyme recognition. Specifically, the strongest association of these nucleotides aligns with previously characterized recognition features for Trm11, requiring G10oU25 pair in the D-arm (Urbonavičius et al., 2006). Our broad sampling across archaeal species supports this finding, reinforcing its conserved functional importance.

Among Euryarchaeotal species, the frequency of predicted m²G10/m²_2_G10 modifications correspond with the expected G10oU25 and U13oU22 (Figure 5A). Moreover, our analysis identifies additional correlations, involving other non-Watson-Crick pairs such as U13oG22 and U13oU22 in Thermococci and Sulfolobales (Additional file: Fig. S8, Table S6).

**Figure 5.**
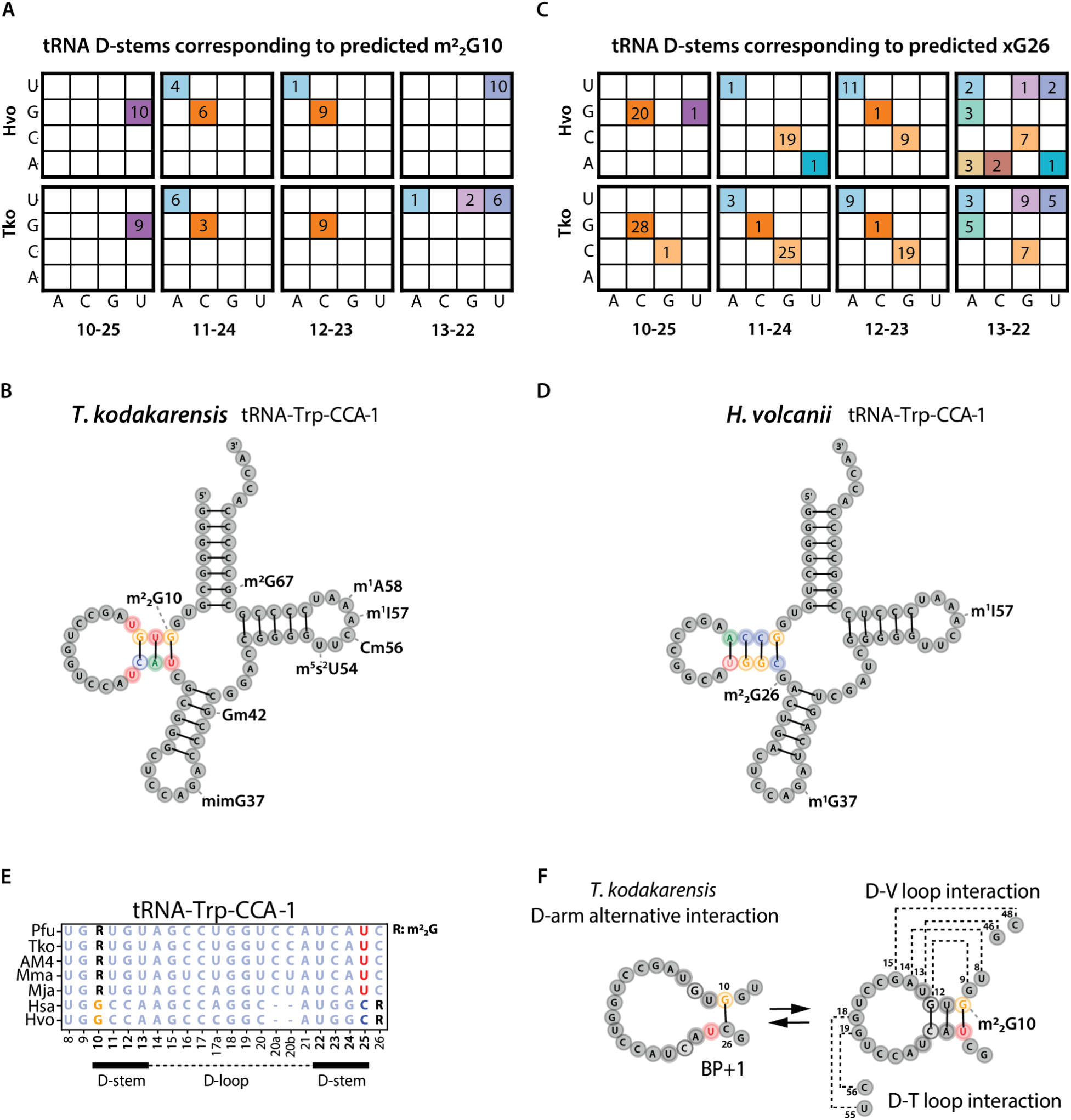
Conserved sequence and structural features correlate with recognition of predicted m^2^ G10 and *x*G26 modified tRNAs. A) Base-pairing identities in D-stems corresponding to high-confidence predictions of m^2^ G10 modifications in *T. kodakarensis* (Tko) and *H. volcanii* (Hvo). Colors represent base-pair combinations and include the number of predicted modified tRNAs for a given pair, such as non-Watson-Crick pairings G10oU25 (purple) and U13oU22 (lavender). B) Predicted secondary structure of tRNA^Trp^ from *T. kodakarensis*, highlighting key structural recognition features for Trm11-mediated m^2^ G10 (G10oU25 and U13oU22). These elements form short, unstable D-arms that facilitate enzyme recognition and modification. C) Base-pair identities in tRNA D-stems associated with high-confidence predictions of *x*G26 modifications, where G10=C25 are conserved and consistent with anti-determinants for m^2^ G10. D) Predicted secondary structure of tRNA^Trp^ from *H. volcanii*, highlighting structural elements and the absence of m^2^ G10 with the formation of a stable G10=C25 W-C pair. E) Sequence alignment of tRNA^Trp^ D-stem regions species, illustrating conserved structural features, particularly G10oU25 versus G10=C25. F) Conceptual illustration of structural instability in the D-arm of tRNA^Trp^ from *T. kodakarensis*. Predicted m^2^ G10 modification stabilizes the functional L-shaped tRNA structure by preventing alternative interactions (e.g. a G10=C26 base pair), promoting proper D-loop and T-loop interactions necessary for canonical folding.

Non-Watson-Crick pairs, particularly GoU pairs, are known to induce structural variations essential for protein binding without necessitating direct protein-base contact (Westhof et al., 2020, 2022). Our extensive cross-clade analysis further supports that these structural variations may contribute to enzyme recognition. Structurally, residues forming non-Watson-Crick pairs result in shorter, less stable D-stems, which often correspond with the presence of predicted m²_2_G10 modifications (Figure 5B & E). Conversely, we observed absences of these modifications in tRNAs possessing more stable D-stems, further supporting the functional role that this modification has in the structural integrity of tRNAs. For instance, within Halobacteria, the tRNA^Trp^ deviates from the conserved G10oU25 pair, instead forming a G10=C25 pair, coinciding with the absence of predicted m²_2_G10 modification (Figure 5C-D). This difference highlights the stringent substrate specificity linked to features like U25, essential for facilitating m²_2_G10-U25 interactions, but not m²_2_G10-C25 interactions.

Similarly, nucleotide features specific to predicted targets of G10oU25 containing tRNAs have been identified as anti-determinants for the action of Trm1, as demonstrated previously in *P. furiosus* (Urbonavičius et al., 2006). Our results support this relationship among the studied Euryarchaeota species, showing a clear association between the presence of G10oU25 and the absence of m^2^ G26 (Figure 5E). Interestingly, Sulfolobales do not exhibit this exclusion pattern and often show co-modification of m²_2_G10 and m²_2_G26. Additionally, Sulfolobales show unique targeting by Trm11 of C12=G23 and A12-U23-containing tRNAs (Additional file: Fig. S8, Table S7). This divergence highlights structural adaptation strategies distinguishing Sulfolobales from other studied archaeal groups. In general, weaker D-arms likely depend on m²_2_G10 to stabilize the tRNA in its functional L-shaped conformation (Steinberg & Cedergren, 1995). In a specific example of the tRNA^Trp^ of *T. kodakarensis*, we see how m²_2_G10 may prevent an alternative interaction between G10=C26 and likely plays a role in most m²_2_G10oU25-containing tRNAs (Figure 5F). Collectively, this multi-clade analysis significantly extends our current knowledge on interplay between tRNA sequence, structure, and modification enzymes.

## Discussion

Comprehensive analyses across diverse archaeal species has revealed distinct modification patterns reflecting adaptive mechanisms shaped by both environmental pressures and phylogenetic constraints. Through interrogating high-throughput RNA sequencing datasets alongside genome protein set analyses, we have expanded the scope of our understanding of post-transcriptional tRNA modifications, uncovering novel relationships, hypothesized environmental adaptations, and extensive new data illuminating the coevolution of tRNA substrates with their modifying enzymes.

One surprising finding is the coordination of m^2^G/m^2^ G at positions 6 and 67 in the acceptor stem of hyperthermophiles, correlating with the presence and conservation of Trm14 orthologs. In contrast, the two Sulfolobales species examined here lack both the coordinated modification at these positions and detectable Trm14 homologs. However, the broader distribution of Trm14 homologs present among Thermoprotei species with optimal growth temperatures exceeding 80℃ suggests that these acceptor stem modifications may extend beyond Euryarchaeota to include hyperthermophilic Sulfolobales. While previous work noted individual modifications at these positions (Yu et al., 2019), our comparative approach clearly indicates their mutual exclusivity, suggesting a previously unrecognized functional interdependence. Despite their exclusive occurrence in hyperthermophiles, these modifications may not play a role in thermal stability, which typically involves T-loop/D-loop interactions at the external globular corner of the L-shape (Grosjean & Oshima, 2007). A prior study (Wolff et al., 2023) suggests m^2^ G6-C67 disrupts the W-C base pair, resulting in a pronounced propeller twist, potentially playing a role in modulating the aminoacylation efficiency of aaRS tRNA charging. With our observations of this coordinated relationship, we hypothesize that as the steric disruptions between m^2^ G6-C67 and C6-m^2^G67 change, charging efficiencies may be regulated for specific tRNAs. Beyond this role, we further speculate that these steric alterations could influence mechanics of translation itself. During translocation, the acceptor stem must navigate steric blocks moving from the A to the P site (Noller et al., 2017). The propeller twist induced by these modifications could impact kinetics and fidelity of this process by altering how the acceptor stem is accommodated. We hypothesize that this could provide hyperthermophiles a strategy to maintain or alter protein synthesis rates in response to environmental shifts, but requires additional study.

Additionally, we identified coordinated modification patterns involving conserved core positions 10 and 26 across multiple archaeal clades. Our results align with, yet also challenge, established models of substrate specificities for Trm1 and Trm11 orthologs. While tRNAs with G10 typically contain a G10oU25 base pair consistent with previous work (Urbonavičius et al., 2006), the co-occurrence of modifications at both position 10 and 26 in different species present a more complex picture. Sulfolobales species, for instance, exhibit m^2^ G10 and m^2^ G26 within the same tRNA^Glu^, despite being previously proposed as anti-determinants in *P. furiosus*, where structural elements required for dimethylation of one site preclude modification at the other.

Broader comparisons allowed us to observe differences in Thermococci, where lower rates of misincorporation at position 10 suggest a predominant pairing of m^2^G10 with m^2^ G26 that either co-occur or are coordinated in the same tRNA, such as in Asn^GTT^. This diversity suggests that while fundamental roles of core modifications are conserved, the enzyme specificities have diverged between archaeal lineages. These modification patterns can be explained by the coevolution of Trm1 and Trm11 with their substrates. For instance, we observed the Trm11 orthologs in Sulfolobales uniquely lack the THUMP domain, which is critical for tRNA binding through recognition of the CCA tail (Hori, 2023). This architectural divergence coincides with the unusual frequency of co-occurring modification patterns as well as variable target substrates. It is plausible that this domain loss is compensated by interactions with accessory proteins, such as Trm112, which is required for Trm11 activity in *H. volcanii* (van Tran et al., 2018). An enzyme complex could possess altered substrate recognition, enabling the bypass of canonical determinants and would otherwise be unsuitable substrates. The unique targeting of tRNAs with C12=G23 or A12oU23 pairs in the D-stem of Sulfolobales further illustrates enzyme and substrate coevolution produce lineage-specific outcomes.

The structural elements governing modification in the D-stem are deeply conserved, though specific implementations vary. In yeast, the formation of m^2^ G26 requires two consecutive GC pairs in the D-stem, whereas bacteria has tolerance for AU pairs (Edqvist et al., 1994; Takeda et al., 2002). Likewise, recent work on human TRMT1 shows it catalyzes either m^2^G or m^2^ G in a substrate-dependent manner governed by base pairs in the D-stem (Xiong et al., 2023). This highlights a conserved structural code for enzyme-substrate recognition that has been fine-tuned throughout evolution. As a general rule, these core modifications have been found to be the key for maintaining the tRNA L-shaped architecture and resistance to degradation (K. Zhang et al., 2025). Methylations, in particular, enhance thermostability by promoting precise H-bonds (e.g. m^1^A favors Hoogsteen pairs, m^2^G/m^2^ G favors GoU pairs), improving base stacking, introducing localized positive charges that interact with the phosphate backbone, and reinforce tertiary interactions (Auffinger & Westhof, 2001; Wolff et al., 2023).

The presence of m^2^ G is especially important, as it can disrupt non-canonical base pairings and prevent alternative, non-functional tRNA conformations. Therefore, the co-occurring modification of both positions 10 and 26 in Sulfolobales may reflect a unique evolutionary strategy to ensure tRNA integrity in environments of high acidity and elevated temperatures.

While our analysis revealed clear nucleotide determinants for some modifications, it also highlighted different classes of enzymes that appear to rely on alternative recognition strategies. For instance, sites predicted to carry m²G6/m²_2_G6, m^1^A9/m^1^G9, and m²G67, do not exhibit strong evidence for associations with specific nucleotide residues within the tRNA body. This aligns with previous biochemical studies on enzymes like Trm10, which suggest substrate preference may be dictated by the overall stability of the folded tRNA rather than local sequence motifs (Strassler et al., 2022). It is plausible that for certain enzymes, recognition depends on global interactions with the L-shaped tertiary structure combined with precise positioning at the catalytic site. This distinction demonstrates the diversity of recognition strategies across modification enzymes and emphasizes that inferring determinants for many modifications requires integrating tertiary structural context.

Distinguishing between these modes of recognition requires accurate and comprehensive modification maps, yet our survey also demonstrates how detection is shaped by the inherent biases of RT-based methods. Modifications that disrupt W-C base pairing, such as m^1^A and m^2^ G, consistently yield high misincorporation signals and are therefore robustly detected. Conversely, modifications like 2’-*O*-methylations (Am, Cm, Gm, Um), which alter the ribose sugar rather than the base-pairing face, cause minimal disruption to reverse transcription and are notoriously difficult to detect with this approach. This limitation warrants caution when interpreting low misincorporation signals, or their absence, as evidence against modification, particularly at conserved sites like Cm56.

These methodological challenges culminate in the classification of many potential modification sites as “Not Determined” (ND), which represent a frontier for discovery. For instance, the misincorporation frequencies at G43 in tRNA^Gln^ of *S. islandicus* likely indicate a novel site of 2’-*O*-methylguanine, but its characterization is beyond the scope of this study. Future work should prioritize validating these ND sites using a combination of orthogonal approaches. Mass spectrometry remains essential for the unambiguous identification of novel modifications. Furthermore, recent advances in direct RNA sequencing, particularly with nanopore technology, offer powerful new tools for mapping modifications on individual, full-length molecules. This technology holds promise for the direct detection of modifications like 2’-*O*-methylations that are challenging to detect using RT-based methods, allowing for a more complete and systematic mapping of these across the archaeal tRNome. Investigating how these newly identified methylations contribute to translational fidelity and stress adaptation in archaea would be a valuable next step.

In summary, this work leverages high-throughput data to map the complex landscape of archaeal tRNA modifications, revealing a dynamic interplay of conserved rules and lineage-specific co-evolution. We present a comprehensive experimental analysis that lays the groundwork for future research by contributing detailed information on tRNA modification profiles, substrate preferences, and evolutionary patterns. These results provide a valuable foundation for guiding targeted biochemical validations, structural studies of enzyme-substrate interactions, and the development of mechanistic models that explain modification specificity and coordination across the domains of life.

## Conclusions

This study provides a comprehensive experimental analysis of archaeal tRNA modifications and the enzymes predicted to install them. By combining high-throughput sequencing with comparative genomic analysis, we uncovered coordinated and clade-specific modification patterns at key tRNA positions and linked these patterns to evolutionary changes in enzyme specificity and domain architecture. Our findings extend the known repertoire of recognition elements and anti-determinants of Trm11 and Trm1 homologs, while also revealing instances where recognition rules are challenged, particularly within Sulfolobales. These results emphasize co-evolution between tRNA structure, modifications, and their enzymes in the context of extreme environmental adaptation. Beyond mapping, this work establishes a framework for future biochemical validation, predictive modeling, and functional studies of tRNA modifications.

## Acknowledgments

We thank Lucas Ferguson and Kathleen Collins for providing reagents for OTTR-seq experiments, and their considerable assistance and consultation in optimizing the protocol for tRNAs. We are grateful to members of the NSF-funded archaeal epitranscriptomics consortium for providing the RNA samples of many species. For *M. maripaludis*, we thank Taiwo Akinyemi, Elliot Shelton, and William B. Whitman, supported by US DOE grant DESC0018028. For *T. kodakarensis* and *T. sp.* AM4, we thank Brett Burkhart. For *S. islandicus*, we thank Rachel Whitaker and Changyi Zhang. For *S. acidocaldarius*, we thank Sonja Albers. For *H. salinarium*, we thank David Crowley. For *M. jannaschii*, we thank Biswarup Mukhopadhyay. We thank Patricia Chan for meaningful comments and edits to the manuscript. This work was supported by funding from the USA National Science Foundation, award #2022065 (to TJS and TML), USA National Institutes of Health, award R35 GM143963 (to TJS), and the USA National Aeronautics and Space Administration, award 80NSSC23K1354 (to TJS).

## Methods

### Culture and RNA isolation

Cultures and total RNA samples for this study were obtained from a combination of in-house growth experiments and collaborations. *H. volcanii,* was cultured in our lab following established protocols (Halohandbook), with three biological replicates that indicated reproducibility for downstream analyses (Additional file: Fig. S9). Cell pellets of *M. maripaludis, M. jannaschii, S. islandicus, S. acidocaldarius,* and *H. salinarum* were kindly provided by the laboratories of William B. Whitman, Bishwarup Mukhopadhyay, Rachel Whitaker, Sonja Albers, and David Crowley, respectively (Crowley et al., 2006; Jäger et al., 2014; Liman et al., 2022; Mukhopadhyay et al., 1999; Tang et al., 2009; Wagner et al., 2012; Whitman et al., 1987; C. Zhang et al., 2013). *T. kodakarensis* and *T. sp.* AM4 were cultured in Tom Santangelo’s lab (Jäger et al., 2014). All RNA extractions from externally provided cell pellets were performed by Tom Santangelo’s lab. Total RNA from cell pellets were extracted as described previously (Tsai et al., 2025). Briefly, cells were treated with BAN reagent (Molecular Research Center, Inc., Cat #BN191). RNA was purified with an equal volume of acid phenol:chloroform:isoamyl alcohol (125:24:1, Thermo Fisher Scientific, Cat #AM9722) and centrifuged to separate the aqueous phase from the organic phase. RNA was then precipitated with 1.5 volumes of isopropanol and 0.1 volume of sodium acetate (Sigma Aldrich, Cat #S7899) at -20C overnight. Finally, the precipitated RNA pellets were washed with 75% ethanol and dissolved in nuclease-free water. Total RNA was used as input for sequencing library construction.

### OTTR-seq library construction

Total RNA was used for generating OTTR-seq libraries, as previously described (Upton et al., 2021). Slight adjustments to the protocol were made to improve tRNA capture. Briefly, total T4 polynucleotide kinase-treated RNA was 3’ tailed using mutant *Bombyx mori* R2 retroelement reverse transcriptase (BoMoC RT) in buffer containing only ddATP for 90 minutes at 30 C°, with the addition of ddGTP for another 30 minutes at 30 C°. This was then heat-inactivated at 65 C° for 5 minutes, and unincorporated ddATP/ddGTP were hydrolyzed by incubation in 5 mM MgCl_2_ and 0.5 units of shrimp alkaline phosphatase (rSAP) at 37 C° for 15 minutes. 5 mM egtazic acid (EGTA) was added and incubated for 100 C° for 5 minutes to stop the reaction and denature GC-rich tRNAs. Reverse transcription was then performed at 37 C° for 30 minutes, followed by heat inactivation at 70 C° for 5 minutes. The remaining RNA and RNA/DNA hybrids were then degraded using 1 unit of RNase A at 50 C° for 10 minutes. cDNA was purified using the MinElute Reaction CleanUp Kit (Qiagen). To reduce adaptor dimers, cDNA was run on a 9% urea-PAGE gel, and tRNA size range (∼60-115bp) was cut out and eluted into gel extraction buffer (300mM NaCl, 10mM Tris; pH 8.0, 1mM EDTA, 0.25% SDS) and concentrated using EtOH precipitation. Size-selected cDNA was then PCR-amplified for 12 cycles using Q5 High-fidelity polymerase (NEB #M0491S) and buffer pack with high GC Enhancer (NEB #B9027S). Amplified libraries were then run on a 6% TBE gel, and tRNA size range (∼60-115bp) was extracted to reduce adaptor dimers further. Gel slices were eluted into a gel extraction buffer (300mM NaCl, 10mM Tris; pH 8.0, 1mM EDTA) followed by concentration using EtOH precipitation. Final libraries were pooled and sequenced for 150 single-end reads on an Illumina NextSeq.

### tRNA Analysis of eXpression (tRAX)

Sequencing adaptors were trimmed from raw reads using cutadapt, v1.18 (Martin, 2011), and read counts were generated for annotated RNAs using tRAX (Holmes et al., 2022). Briefly, trimmed reads were mapped to their respective reference databases: *Methanococcus maripaludis* S2, *Methanocaldococcus jannaschii* DSM 2661, *Sulfolobus islandicus* M.16.4, *Sulfolobus acidocaldarius* DSM 639, *Thermococcus kodakarensis* KOD1, *Thermococcus* sp. AM4, *Pyrococcus furiosus* DSM 3638, *Halobacterium salinarum* R1, and *Haloferax volcanii* DS2.

Each reference tRAX database includes both the genome sequences and the mature tRNA annotations obtained from Data Release 22 of the Genomic tRNA Database (GtRNAdb) (Chan & Lowe, 2016). Trimmed reads were mapped using Bowtie2 (Langmead & Salzberg, 2012) in very-sensitive mode with the following parameters to allow for a maximum of 100 alignments per read: –very-sensitive –ignore-quals –np 5 -k 100. Mapped reads were then filtered to retain only the “best mapping” alignments. Raw read counts of tRNAs and other small RNA types were computed using tRNA annotations from GtRNAdb, and annotations from NCBI RefSeq (Goldfarb et al., 2025). Raw read counts were then normalized using DESeq2 (Love et al., 2014).

### tRNA modification analysis

Sites of known tRNA modifications were compiled from Modomics (Cappannini et al., 2023) and relevant literature (Hirata et al., 2019; Krishnamohan et al., 2019; Wolff et al., 2023; Yu et al., 2019). Modification types for *M. maripaludis*, *M. jannaschii*, *S. acidocaldarius*, *T. kodakarensis, P. furiosus*, and *H. volcanii* were curated in a tab-delimited table (Additional File: Table S0) and mapped to their corresponding tRNA sequence alignments. All tRNA nucleotide positions presented in this work refer to Sprinzl positions, a standard numbering system for tRNAs that assigns consistent numbers to conserved positions across all tRNAs (Sprinzl et al., 1998). Multiple naming conventions are used to refer to tRNAs which include tDNA locus name and transcript name with superscript anticodon, switching from T → U, however both are equivalent. Position-specific MI frequencies were stratified by nucleotide identity for known modifications and a set of bases classified as ND (Not Determined). ND includes positions where there are currently no published annotations for modification in the specific species studied. This group likely contains a mixture of both modified and unmodified bases, but no prior data exists to support a specific classification. ND serves as a background reference for MI behavior in the absence of modification expectations. Known modification positions were mapped onto orthologous tRNAs in unannotated species within phylogenetic clades, focusing on sites with conserved reference nucleotides and shared rates of misincorporation.

To assess whether observed misincorporation frequencies were statistically significant relative to background, we applied tMAP (K. Zhang et al., 2025) using aligned tRNA reads from tRAX (Holmes et al., 2022) as input. At each position misincorporation frequency was calculated as the number of non-reference base calls divided by total read coverage. A beta distribution was modeled using these empirical counts, incorporating pseudocounts derived from the mean and variance of misincorporation rates at known modified positions. A modification threshold was empirically determined as the 75th percentile of misincorporation rates, stratified by reference base identity. P-values were computed using the cumulative distribution function of the beta distribution, representing the probability of observing a misincorporation frequency equal to or greater than the observed value under the null model.

Modification detection relied on two key metrics: the observed misincorporation rate at each position, and the statistical significance (adjusted p-values) of these observed rates. We implemented a base-specific thresholding strategy that accounts for background misincorporation rates inherent to each base. Specifically, cytosine and guanine-derived positions tend to exhibit elevated background rates (often exceeding 5%), resulting in statistically significant p-values despite the probable absence of true modification (Additional File: Table S2). To mitigate false-positive predictions from these elevated baseline rates, we adopted higher confidence thresholds for these bases.

Sites of predicted modification were classified as high-confidence if the MI frequency exceeded 10% for G/C bases or 5% for A/U bases, and had statistically significant adjusted p-values (<0.05). Positions with significant p-values but MI rates below these base-specific threshold were classified as moderate-confidence sites due to the observed mixture of true and false positives in our training set. This classification strategy maintains sensitivity for modifications capable of inducing reverse transcription (RT) misincorporation while enhancing specificity across variable signal contexts.

Variation in modification detection across species, modification types, and positions primarily reflected differences in MI rate distribution within distinct sequence contexts and tRNA positions. Therefore, statistical significance at similar MI rates varied depending on how closely a given rate aligned with its background distribution. Although low read coverage occasionally limited detection sensitivity, the primary determinants of statistical significance were the depth and magnitude of MI at each position.

In instances where multiple modification types are known to occur at the same position, relative misincorporation frequency was used to infer the likely modification. For example, m^2^ G generally causes higher misincorporation than m^2^G at position 10, enabling us to discriminate between these two forms in specific archaeal species. This distinction was also partially observable at positions 6 and 26, though with reduced confidence. In some instances, misincorporation patterns were not sufficiently distinct to confidently infer modification type. In such cases, known modification types from closely related species with identical reference bases were used to assign a most-likely modification classification.

### Homolog search and domain analysis

To search for archaeal homologs and compare domain architectures, we collected representative tRNA modification enzymes that were previously characterized (Constantinesco et al., 1998; Jackman et al., 2003; Menezes et al., 2011; Urbonavičius et al., 2006). Each representative was queried across 220 archaeal genomes included in GtRNAdb release 22 (Chan & Lowe, 2016) using BLASTP (Altschul et al., 1997) with default parameters, then results were filtered to select hits with the lowest e-value for each species and representative. Domain analysis and annotation of highest scoring hits was performed using the InterProScan software package (v5.65-97.0) (Jones et al., 2014) with the FunFam (4.3.0), MobiDBLite (v2.0), NCBIfam (v13.0), PANTHER (v18.0), SuperFamily (v1.75), CDD (v3.20), Pfam (v36.0), SMART (v9.0), PRINTS (v42.0) and PIRSF (v2023_05) databases. As e-values are specific to each InterPro database and each utilizes their own e-value post-processing, Pfam (Mistry et al., 2021) and CDD (Lu et al., 2020) were used preferably when domains were found. All matches were considered tentative hits.

### Phylogenetic analysis

The evolutionary relationships among each of the tRNA modification enzymes were inferred using maximum likelihood phylogenies derived from clustered BlastP hits of representatives and their subsequent multiple sequence alignments. In brief, hits for each representative were clustered using MMseq2 (Steinegger & Söding, 2017) using the parameters --min-seq-id 0.3 --cov-mode 0. Clustered groups were subset and aligned with MAFFT (Rozewicki et al., 2019) using default parameters, then concatenated and aligned again using the parameters --retree 1 --maxiterate 0.

All maximum likelihood phylogenetic trees were estimated using IQ-TREE 2 (v2.1.0) (Nguyen et al., 2015). Selection of the best-fit model of amino acid substitution was inferred for the phylogenies using the ModelFinder function in IQ-TREE (LG+F+R6). Custom python scripts were used to prune trees by keeping only BlastP hits of each representative with the lowest e-value in a genome. Trees of each representative and its best-scoring homologs were plotted using the Interactive Tree Of Life (Letunic & Bork, 2021). Predicted domain architectures were annotated onto the phylogenies.

### tRNA secondary structure

The R2DT software (McCann et al., 2025) was used to predict and illustrate tRNA secondary structures. Predicted modifications and tertiary interactions were illustrated on secondary structure images manually.

## Notes

### Competing Interest Statement

The authors have declared no competing interest.

### Summary of Updates

Text revisions in all sections; Figure color changes; Supplemental Figure legends revised

## References

1. Altschul, S. F., Madden, T. L., Schäffer, A. A., Zhang, J., Zhang, Z., Miller, W., & Lipman, D. J. (1997). Gapped BLAST and PSI-BLAST: A new generation of protein database search programs. Nucleic Acids Research, 25(17), 3389–3402. 10.1093/nar/25.17.3389

2. Auffinger, P., & Westhof, E. (2001). An extended structural signature for the tRNA anticodon loop. RNA (New York, N.Y.), 7(3), 334–341. 10.1017/s1355838201002382

3. Behrens, A., Rodschinka, G., & Nedialkova, D. D. (2021). High-resolution quantitative profiling of tRNA abundance and modification status in eukaryotes by mim-tRNAseq. Molecular Cell, 81(8), 1802–1815.e7. 10.1016/j.molcel.2021.01.028

4. Bujnicki, J. M., Feder, M., Radlinska, M., & Blumenthal, R. M. (2002). Structure Prediction and Phylogenetic Analysis of a Functionally Diverse Family of Proteins Homologous to the MT-A70 Subunit of the Human mRNA:m6A Methyltransferase. Journal of Molecular Evolution, 55(4), 431–444. 10.1007/s00239-002-2339-8

5. Cappannini, A., Ray, A., Purta, E., Mukherjee, S., Boccaletto, P., Moafinejad, S. N., Lechner, A., Barchet, C., Klaholz, B. P., Stefaniak, F., & Bujnicki, J. M. (2023). MODOMICS: A database of RNA modifications and related information. 2023 update. Nucleic Acids Research, gkad1083. 10.1093/nar/gkad1083

6. Chan, P. P., & Lowe, T. M. (2016). GtRNAdb 2.0: An expanded database of transfer RNA genes identified in complete and draft genomes. Nucleic Acids Research, 44(D1), D184–D189. 10.1093/nar/gkv1309

7. Constantinesco, F., Benachenhou, N., Motorin, Y., & Grosjean, H. (1998). The tRNA(guanine-26,N2-N2) methyltransferase (Trm1) from the hyperthermophilic archaeon Pyrococcus furiosus: Cloning, sequencing of the gene and its expression in Escherichia coli. Nucleic Acids Research, 26(16), 3753–3761.

8. Cozen, A. E., Quartley, E., Holmes, A. D., Hrabeta-Robinson, E., Phizicky, E. M., & Lowe, T. M. (2015). ARM-seq: AlkB-facilitated RNA methylation sequencing reveals a complex landscape of modified tRNA fragments. Nature Methods, 12(9), 879–884. 10.1038/nmeth.3508

9. Crowley, D. J., Boubriak, I., Berquist, B. R., Clark, M., Richard, E., Sullivan, L., DasSarma, S., & McCready, S. (2006). The uvrA, uvrB and uvrC genes are required for repair of ultraviolet light induced DNA photoproducts in Halobacterium sp. NRC-1. Saline Systems, 2, 11. 10.1186/1746-1448-2-11

10. de Crecy-Lagard, V., Brochier-Armanet, C., Urbonavičius, J., Fernandez, B., Phillips, G., Lyons, B., Noma, A., Alvarez, S., Droogmans, L., Armengaud, J., & Grosjean, H. (2010). Biosynthesis of Wyosine Derivatives in tRNA: An Ancient and Highly Diverse Pathway in Archaea. 16.

11. de Crécy-Lagard, V., Boccaletto, P., Mangleburg, C. G., Sharma, P., Lowe, T. M., Leidel, S. A., & Bujnicki, J. M. (2019). Matching tRNA modifications in humans to their known and predicted enzymes. Nucleic Acids Research, 47(5), 2143–2159. 10.1093/nar/gkz011

12. Edqvist, J., Blomqvist, K., & Straaby, K. (1994). Structural Elements in Yeast tRNAs Required for Homologous Modification of Guanosine-26 into Dimethylguanosine-26 by the Yeast Trm1 tRNA-Modifying Enzyme. Biochemistry, 33(32), 9546–9551. 10.1021/bi00198a021

13. Erber, L., Hoffmann, A., Fallmann, J., Betat, H., Stadler, P. F., & Mörl, M. (2020). LOTTE-seq (Long hairpin oligonucleotide based tRNA high-throughput sequencing): Specific selection of tRNAs with 3’-CCA end for high-throughput sequencing. RNA Biology, 17(1), 23–32. 10.1080/15476286.2019.1664250

14. Fislage, M., Roovers, M., Tuszynska, I., Bujnicki, J. M., Droogmans, L., & Versées, W. (2012). Crystal structures of the tRNA:m 2 G6 methyltransferase Trm14/TrmN from two domains of life. Nucleic Acids Research, 40(11), 5149–5161. 10.1093/nar/gks163

15. Fluke, K. A., Fuchs, R. T., Tsai, Y.-L., Talbott, V., Elkins, L., Febvre, H. P., Dai, N., Wolf, E. J., Burkhart, B. W., Schiltz, J., Brett Robb, G., Corrêa, I. R., & Santangelo, T. J. (2024). The extensive m5C epitranscriptome of Thermococcus kodakarensis is generated by a suite of RNA methyltransferases that support thermophily. Nature Communications, 15(1), 7272. 10.1038/s41467-024-51410-w

16. Gabant, G., Auxilien, S., Tuszynska, I., Locard, M., Gajda, M. J., Chaussinand, G., Fernandez, B., Dedieu, A., Grosjean, H., Golinelli-Pimpaneau, B., Bujnicki, J. M., & Armengaud, J. (2006). THUMP from archaeal tRNA:m22G10 methyltransferase, a genuine autonomously folding domain. Nucleic Acids Research, 34(9), 2483–2494. 10.1093/nar/gkl145

17. Goldfarb, T., Kodali, V. K., Pujar, S., Brover, V., Robbertse, B., Farrell, C. M., Oh, D.-H., Astashyn, A., Ermolaeva, O., Haddad, D., Hlavina, W., Hoffman, J., Jackson, J. D., Joardar, V. S., Kristensen, D., Masterson, P., McGarvey, K. M., McVeigh, R., Mozes, E., … Murphy, T. D. (2025). NCBI RefSeq: Reference sequence standards through 25 years of curation and annotation. Nucleic Acids Research, 53(D1), D243–D257. 10.1093/nar/gkae1038

18. Grosjean, H. (2010). Deciphering synonymous codons in the three domains of life: Co-evolution with specific tRNA modification enzymes. FEBS Letters, 13.

19. Grosjean, H., Gaspin, C., Marck, C., Decatur, W. A., & de Crécy-Lagard, V. (2008). RNomics and Modomics in the halophilic archaea Haloferax volcanii: Identification of RNA modification genes. BMC Genomics, 26.

20. Grosjean, H., & Oshima, T. (2007). How Nucleic Acids Cope with High Temperature. In Physiology and Biochemistry of Extremophiles (pp. 39–56). John Wiley & Sons, Ltd. 10.1128/9781555815813.ch4

21. Gupta, R. (1984). Halobacterium volcanii tRNAs. Identification of 41 tRNAs covering all amino acids, and the sequences of 33 class I tRNAs. Journal of Biological Chemistry, 259(15), 9461–9471. 10.1016/S0021-9258(17)42723-2

22. Gustafsson, H. T., Galan, C., Yu, T., Upton, H. E., Ferguson, L., Kaymak, E., Weng, Z., Collins, K., & Rando, O. J. (2022). *Deep sequencing of yeast and mouse tRNAs and tRNA fragments using OTTR* [Preprint]. Genomics. 10.1101/2022.02.04.479139

23. Helm, M. (2006). Post-transcriptional nucleotide modification and alternative folding of RNA. Nucleic Acids Research, 34(2), 721. 10.1093/nar/gkj471

24. Hernandez-Alias, X., Katanski, C. D., Zhang, W., Assari, M., Watkins, C. P., Schaefer, M. H., Serrano, L., & Pan, T. (2022). Single-read tRNA-seq analysis reveals coordination of tRNA modification and aminoacylation and fragmentation. Nucleic Acids Research, gkac1185. 10.1093/nar/gkac1185

25. Hirata, A., Nishiyama, S., Tamura, T., Yamauchi, A., & Hori, H. (2016). Structural and functional analyses of the archaeal tRNA m ^2^ G/m ^2^ _2_ G10 methyltransferase aTrm11 provide mechanistic insights into site specificity of a tRNA methyltransferase that contains common RNA-binding modules. Nucleic Acids Research, 44(13), 6377–6390. 10.1093/nar/gkw561

26. Hirata, A., Suzuki, T., Nagano, T., Fujii, D., Okamoto, M., Sora, M., Lowe, T. M., Kanai, T., Atomi, H., Suzuki, T., & Hori, H. (2019). Distinct Modified Nucleosides in tRNA ^Trp^ from the Hyperthermophilic Archaeon *Thermococcus kodakarensis* and Requirement of tRNA m ^2^ G10/m ^2^ _2_ G10 Methyltransferase (Archaeal Trm11) for Survival at High Temperatures. Journal of Bacteriology, 201(21), e00448–19,/jb/201/21/JB.00448-19.atom. 10.1128/JB.00448-19

27. Höfer, K., & Jäschke, A. (2018). Epitranscriptomics: RNA Modifications in Bacteria and Archaea. Microbiology Spectrum, 6(3), 10.1128/microbiolspec.rwr-0015–2017. 10.1128/microbiolspec.rwr-0015-2017

28. Holmes, A. D., Howard, J. M., Chan, P. P., & Lowe, T. M. (2022). tRNA Analysis of eXpression (tRAX): A tool for integrating analysis of tRNAs, tRNA-derived small RNAs, and tRNA modifications (p. 2022.07.02.498565). bioRxiv. 10.1101/2022.07.02.498565

29. Hori, H. (2014). Methylated nucleosides in tRNA and tRNA methyltransferases. Frontiers in Genetics, 26.

30. Hori, H. (2019). Regulatory Factors for tRNA Modifications in Extreme-Thermophilic Bacterium Thermus thermophilus. Frontiers in Genetics, 10, 204. 10.3389/fgene.2019.00204

31. Hori, H. (2023). Transfer RNA Modification Enzymes with a Thiouridine Synthetase, Methyltransferase and Pseudouridine Synthase (THUMP) Domain and the Nucleosides They Produce in tRNA. Genes, 14(2), 382. 10.3390/genes14020382

32. Hori, H., Kawamura, T., Awai, T., Ochi, A., Yamagami, R., Tomikawa, C., & Hirata, A. (2018). Transfer RNA Modification Enzymes from Thermophiles and Their Modified Nucleosides in tRNA. Microorganisms, 6(4), 110. 10.3390/microorganisms6040110

33. Ihsanawati Nishimoto, M., Higashijima, K., Shirouzu, M., Grosjean, H., Bessho, Y., & Yokoyama, S. (2008). Crystal Structure of tRNA *N*2,*N*2-Guanosine Dimethyltransferase Trm1 from *Pyrococcus horikoshii*. Journal of Molecular Biology, 383(4), 871–884. 10.1016/j.jmb.2008.08.068

34. Jackman, J. E., & Alfonzo, J. D. (2013). Transfer RNA modifications: Nature’s combinatorial chemistry playground: Transfer RNA modifications. Wiley Interdisciplinary Reviews: RNA, 4(1), 35–48. 10.1002/wrna.1144

35. Jackman, J. E., Montange, R. K., Malik, H. S., & Phizicky, E. M. (2003). Identification of the yeast gene encoding the tRNA m1G methyltransferase responsible for modification at position 9. RNA (New York, N.Y.), 9(5), 574–585. 10.1261/rna.5070303

36. Jäger, D., Förstner, K. U., Sharma, C. M., Santangelo, T. J., & Reeve, J. N. (2014). Primary transcriptome map of the hyperthermophilic archaeon Thermococcus kodakarensis. BMC Genomics, 15(1), 684. 10.1186/1471-2164-15-684

37. Jones, P., Binns, D., Chang, H.-Y., Fraser, M., Li, W., McAnulla, C., McWilliam, H., Maslen, J., Mitchell, A., Nuka, G., Pesseat, S., Quinn, A. F., Sangrador-Vegas, A., Scheremetjew, M., Yong, S.-Y., Lopez, R., & Hunter, S. (2014). InterProScan 5: Genome-scale protein function classification. Bioinformatics, 30(9), 1236–1240. 10.1093/bioinformatics/btu031

38. Kimura, S., Dedon, P. C., & Waldor, M. K. (2020). Comparative tRNA sequencing and RNA mass spectrometry for surveying tRNA modifications. Nature Chemical Biology, 16(9), 964–972. 10.1038/s41589-020-0558-1

39. Kowalak, J. A., Dalluge, J. J., McCloskey, J. A., & Stetter, K. O. (1994). The Role of Posttranscriptional Modification in Stabilization of Transfer RNA from Hyperthermophiles. Biochemistry, 33(25), 7869–7876. 10.1021/bi00191a014

40. Krishnamohan, A., Dodbele, S., & Jackman, J. (2019). Insights into Catalytic and tRNA Recognition Mechanism of the Dual-Specific tRNA Methyltransferase from Thermococcus kodakarensis. Genes, 10(2), 100. 10.3390/genes10020100

41. Krishnamohan, A., & Jackman, J. E. (2019). A Family Divided: Distinct Structural and Mechanistic Features of the SpoU-TrmD (SPOUT) Methyltransferase Superfamily. Biochemistry, 58(5), 336–345. 10.1021/acs.biochem.8b01047

42. Langmead, B., & Salzberg, S. L. (2012). Fast gapped-read alignment with Bowtie 2. Nature Methods, 9(4), 357–359. 10.1038/nmeth.1923

43. Letunic, I., & Bork, P. (2021). Interactive Tree Of Life (iTOL) v5: An online tool for phylogenetic tree display and annotation. Nucleic Acids Research, 49(W1), W293–W296. 10.1093/nar/gkab301

44. Li, J., Li, H., Long, T., Dong, H., Wang, E.-D., & Liu, R.-J. (2019). Archaeal NSUN6 catalyzes m5C72 modification on a wide-range of specific tRNAs. Nucleic Acids Research, 47(4), 2041–2055. 10.1093/nar/gky1236

45. Liman, G. L. S., Stettler, M. E., & Santangelo, T. J. (2022). Transformation Techniques for the Anaerobic Hyperthermophile Thermococcus kodakarensis. Methods in Molecular Biology (Clifton, N.J.), 2522, 87–104. 10.1007/978-1-0716-2445-6_5

46. Lorenz, C., Lünse, C., & Mörl, M. (2017). tRNA Modifications: Impact on Structure and Thermal Adaptation. Biomolecules, 7(4), 35. 10.3390/biom7020035

47. Love, M. I., Huber, W., & Anders, S. (2014). Moderated estimation of fold change and dispersion for RNA-seq data with DESeq2. Genome Biology, 15(12), 550. 10.1186/s13059-014-0550-8

48. Lu, S., Wang, J., Chitsaz, F., Derbyshire, M. K., Geer, R. C., Gonzales, N. R., Gwadz, M., Hurwitz, D. I., Marchler, G. H., Song, J. S., Thanki, N., Yamashita, R. A., Yang, M., Zhang, D., Zheng, C., Lanczycki, C. J., & Marchler-Bauer, A. (2020). CDD/SPARCLE: The conserved domain database in 2020. Nucleic Acids Research, 48(D1), D265–D268. 10.1093/nar/gkz991

49. Martin, M. (2011). Cutadapt removes adapter sequences from high-throughput sequencing reads. EMBnet.Journal, 17(1), Article 1. 10.14806/ej.17.1.200

50. McCann, H., Meade, C. D., Williams, L. D., Petrov, A. S., Johnson, P. Z., Simon, A. E., Hoksza, D., Nawrocki, E. P., Chan, P. P., Lowe, T. M., Ribas, C. E., Sweeney, B. A., Madeira, F., Anyango, S., Appasamy, S. D., Deshpande, M., Varadi, M., Velankar, S., Zirbel, C. L., … Petrov, A. I. (2025). R2DT: A comprehensive platform for visualizing RNA secondary structure. Nucleic Acids Research, 53(4), gkaf032. 10.1093/nar/gkaf032

51. Menezes, S., Gaston, K. W., Krivos, K. L., Apolinario, E. E., Reich, N. O., Sowers, K. R., Limbach, P. A., & Perona, J. J. (2011). Formation of m2G6 in Methanocaldococcus jannaschii tRNA catalyzed by the novel methyltransferase Trm14. Nucleic Acids Research, 39(17), 7641–7655. 10.1093/nar/gkr475

52. Mistry, J., Chuguransky, S., Williams, L., Qureshi, M., Salazar, G. A., Sonnhammer, E. L. L., Tosatto, S. C. E., Paladin, L., Raj, S., Richardson, L. J., Finn, R. D., & Bateman, A. (2021). Pfam: The protein families database in 2021. Nucleic Acids Research, 49(D1), D412–D419. 10.1093/nar/gkaa913

53. Motorin, Y., & Helm, M. (2011). RNA nucleotide methylation. Wiley Interdisciplinary Reviews. RNA, 2(5), 611–631. 10.1002/wrna.79

54. Mukhopadhyay, B., Johnson, E. F., & Wolfe, R. S. (1999). Reactor-Scale Cultivation of the Hyperthermophilic Methanarchaeon Methanococcus jannaschii to High Cell Densities. Applied and Environmental Microbiology, 65(11), 5059–5065.

55. Nakano, Y., Gamper, H., McGuigan, H., Maharjan, S., Li, J., Sun, Z., Yigit, E., Grünberg, S., Krishnan, K., Li, N.-S., Piccirilli, J. A., Kleiner, R., Nichols, N., Gregory, B. D., & Hou, Y.-M. (2025). Genome-wide profiling of tRNA modifications by Induro-tRNAseq reveals coordinated changes. Nature Communications, 16(1), 1047. 10.1038/s41467-025-56348-1

56. Nguyen, L.-T., Schmidt, H. A., von Haeseler, A., & Minh, B. Q. (2015). IQ-TREE: A Fast and Effective Stochastic Algorithm for Estimating Maximum-Likelihood Phylogenies. Molecular Biology and Evolution, 32(1), 268–274. 10.1093/molbev/msu300

57. Noller, H. F., Lancaster, L., Zhou, J., & Mohan, S. (2017). The ribosome moves: RNA mechanics and translocation. Nature Structural & Molecular Biology, 24(12), 1021–1027. 10.1038/nsmb.3505

58. Ohira, T., & Suzuki, T. (2024). Transfer RNA modifications and cellular thermotolerance. Molecular Cell, 84(1), 94–106. 10.1016/j.molcel.2023.11.041

59. Ontiveros, R. J., Stoute, J., & Liu, K. F. (2019). The chemical diversity of RNA modifications. The Biochemical Journal, 476(8), 1227–1245. 10.1042/BCJ20180445

60. Reno, M. L., Held, N. L., Fields, C. J., Burke, P. V., & Whitaker, R. J. (2009). Biogeography of the Sulfolobus islandicus pan-genome. Proceedings of the National Academy of Sciences, 106(21), 8605–8610. 10.1073/pnas.0808945106

61. Roovers, M., Droogmans, L., & Grosjean, H. (2021). Post-Transcriptional Modifications of Conserved Nucleotides in the T-Loop of tRNA: A Tale of Functional Convergent Evolution. Genes, 12(2), 140. 10.3390/genes12020140

62. Rozewicki, J., Li, S., Amada, K. M., Standley, D. M., & Katoh, K. (2019). MAFFT-DASH: Integrated protein sequence and structural alignment. Nucleic Acids Research, 47(W1), W5–W10. 10.1093/nar/gkz342

63. Sas-Chen, A., Thomas, J. M., Matzov, D., Taoka, M., Nance, K. D., Nir, R., Bryson, K. M., Shachar, R., Liman, G. L. S., Burkhart, B. W., Gamage, S. T., Nobe, Y., Briney, C. A., Levy, M. J., Fuchs, R. T., Robb, G. B., Hartmann, J., Sharma, S., Lin, Q., … Schwartz, S. (2020). Dynamic RNA acetylation revealed by quantitative cross-evolutionary mapping. Nature, 583(7817), 638–643. 10.1038/s41586-020-2418-2

64. Schicho, R. N., Ma, K., Adams, M. W., & Kelly, R. M. (1993). Bioenergetics of sulfur reduction in the hyperthermophilic archaeon Pyrococcus furiosus. Journal of Bacteriology, 175(6), 1823–1830. 10.1128/jb.175.6.1823-1830.1993

65. Schultz, S. K., & Kothe, U. (2024). RNA modifying enzymes shape tRNA biogenesis and function. Journal of Biological Chemistry, 107488. 10.1016/j.jbc.2024.107488

66. Shigi, N. (2018). Recent Advances in Our Understanding of the Biosynthesis of Sulfur Modifications in tRNAs. Frontiers in Microbiology, 9, 2679. 10.3389/fmicb.2018.02679

67. Shigi, N., Suzuki, T., Terada, T., Shirouzu, M., Yokoyama, S., & Watanabe, K. (2006). Temperature-dependent Biosynthesis of 2-Thioribothymidine of Thermus thermophilus tRNA. Journal of Biological Chemistry, 281(4), 2104–2113. 10.1074/jbc.M510771200

68. Sokołowski, M., Klassen, R., Bruch, A., Schaffrath, R., & Glatt, S. (2018). Cooperativity between different tRNA modifications and their modification pathways. Biochimica Et Biophysica Acta. Gene Regulatory Mechanisms, 1861(4), 409–418. 10.1016/j.bbagrm.2017.12.003

69. Sprinzl, M., Horn, C., Brown, M., Ioudovitch, A., & Steinberg, S. (1998). Compilation of tRNA sequences and sequences of tRNA genes. Nucleic Acids Research, 26(1), 148–153. 10.1093/nar/26.1.148

70. Steinberg, S., & Cedergren, R. (1995). A correlation between N2-dimethylguanosine presence and alternate tRNA conformers. RNA (New York, N.Y.), 1(9), 886–891.

71. Steinegger, M., & Söding, J. (2017). MMseqs2 enables sensitive protein sequence searching for the analysis of massive data sets. Nature Biotechnology, 35(11), Article 11. 10.1038/nbt.3988

72. Strassler, S. E., Bowles, I. E., Dey, D., Jackman, J. E., & Conn, G. L. (2022). Tied up in knots: Untangling substrate recognition by the SPOUT methyltransferases. Journal of Biological Chemistry, 298(10), 102393. 10.1016/j.jbc.2022.102393

73. Takeda, H., Hori, H., & Endo, Y. (2002). Identification of Aquifex aeolicus tRNA (m22G26) methyltransferase gene. Nucleic Acids Symposium Series, 2(1), 229–230. 10.1093/nass/2.1.229

74. Tang, S.-L., Tarasov, V., Athanasopoulos, V., Bath, C., Wendoloski, D., Ferrer, C., Pfeiffer, M., Pohlschroder, M., Santos, F., Allers, T., Eichler, J., Camakaris, H., McAlpine, T., Burns, D., Porter, K., Cukalac, T., & Russ, B. (2009). Melissa Holmes Masahiro Kamekura Wan Lam Stewart Nuttall Wayne Woods Peter Jablonski Juan Serrano Katrina Ngui Josefa Antón.

75. Tsai, Y.-L., Wolf, E. J., Fluke, K. A., Fuchs, R. T., Dai, N., Johnson, S. R., Sun, Z., Elkins, L., Burkhart, B. W., Santangelo, T. J., & Corrêa, I. R. (2025). Comprehensive nucleoside analysis of archaeal RNA modification profiles reveals an m7G in the conserved P loop of 23S rRNA. Cell Reports, 44(4), 115471. 10.1016/j.celrep.2025.115471

76. Upton, H. E., Ferguson, L., Temoche-Diaz, M. M., Liu, X., Pimentel, S. C., Ingolia, N. T., Schekman, R., & Collins, K. (2021). *Low-bias ncRNA Libraries using Ordered Two-Template Relay: Serial Template Jumping by a Modified Retroelement Reverse Transcriptase* [Preprint]. Biochemistry. 10.1101/2021.04.30.442027

77. Urbonavičius, J., Armengaud, J., & Grosjean, H. (2006). Identity Elements Required for Enzymatic Formation of *N*2,*N*2-dimethylguanosine from *N*2-monomethylated Derivative and its Possible Role in Avoiding Alternative Conformations in Archaeal tRNA. Journal of Molecular Biology, 357(2), 387–399. 10.1016/j.jmb.2005.12.087

78. van Tran, N., Muller, L., Ross, R. L., Lestini, R., Létoquart, J., Ulryck, N., Limbach, P. A., de Crécy-Lagard, V., Cianférani, S., & Graille, M. (2018). Evolutionary insights into Trm112-methyltransferase holoenzymes involved in translation between archaea and eukaryotes. Nucleic Acids Research, 46(16), 8483–8499. 10.1093/nar/gky638

79. Wagner, M., van Wolferen, M., Wagner, A., Lassak, K., Meyer, B. H., Reimann, J., & Albers, S.-V. (2012). Versatile Genetic Tool Box for the Crenarchaeote Sulfolobus acidocaldarius. Frontiers in Microbiology, 3, 214. 10.3389/fmicb.2012.00214

80. Westhof, E., Liang, S., Tong, X., Ding, X., Zheng, L., & Dai, F. (2020). Unusual tertiary pairs in eukaryotic tRNAAla. RNA, 26(11), 1519. 10.1261/rna.076299.120

81. Westhof, E., Thornlow, B., Chan, P. P., & Lowe, T. M. (2022). Eukaryotic tRNA sequences present conserved and amino acid-specific structural signatures. Nucleic Acids Research, 50(7), 4100–4112. 10.1093/nar/gkac222

82. Whitman, W. B., Sohn, S., Kuk, S., & Xing, R. (1987). Role of Amino Acids and Vitamins in Nutrition of Mesophilic Methanococcus spp. Applied and Environmental Microbiology, 53(10), 2373–2378.

83. Wolff, P., Lechner, A., Droogmans, L., Grosjean, H., & Westhof, E. (2023). Identification of Up47 in three thermophilic archaea, one mesophilic archaeon and one hyperthermophilic bacterium. RNA, rna.079546.122. 10.1261/rna.079546.122

84. Xiong, Q.-P., Li, J., Li, H., Huang, Z.-X., Dong, H., Wang, E.-D., & Liu, R.-J. (2023). Human TRMT1 catalyzes m2G or m22G formation on tRNAs in a substrate-dependent manner. Science China. Life Sciences. 10.1007/s11427-022-2295-0

85. Yared, M.-J., Marcelot, A., & Barraud, P. (2024). Beyond the Anticodon: tRNA Core Modifications and Their Impact on Structure, Translation and Stress Adaptation. Genes, 15(3), Article 3. 10.3390/genes15030374

86. Yu, N., Jora, M., Solivio, B., Thakur, P., Acevedo-Rocha, C. G., Randau, L., de Crécy-Lagard, V., Addepalli, B., & Limbach, P. A. (2019). tRNA Modification Profiles and Codon-Decoding Strategies in *Methanocaldococcus jannaschii*. Journal of Bacteriology, 201(9), e00690–18, /jb/201/9/JB.00690-18.atom. 10.1128/JB.00690-18

87. Zhang, C., Cooper, T. E., Krause, D. J., & Whitaker, R. J. (2013). Augmenting the Genetic Toolbox for Sulfolobus islandicus with a Stringent Positive Selectable Marker for Agmatine Prototrophy. Applied and Environmental Microbiology, 79(18), 5539–5549. 10.1128/AEM.01608-13

88. Zhang, K., Manning, A. C., Lentini, J. M., Howard, J., Dalwigk, F., Maroofian, R., Efthymiou, S., Chan, P., Eliseev, S. I., Yang, Z., Chang, H., Karimiani, E. G., Bakhshoodeh, B., Houlden, H., Kaiser, S. M., Lowe, T. M., & Fu, D. (2025). Human TRMT1 and TRMT1L paralogs ensure the proper modification state, stability, and function of tRNAs. Cell Reports, 44(1), 115092. 10.1016/j.celrep.2024.115092

89. Zhang, W., & Westhof, E. (2025). The Biology of tRNA t6A Modification and Hypermodifications—Biogenesis and Disease Relevance. Journal of Molecular Biology, 169091. 10.1016/j.jmb.2025.169091

90. Zheng, G., Qin, Y., Clark, W. C., Dai, Q., Yi, C., He, C., Lambowitz, A. M., & Pan, T. (2015). Efficient and quantitative high-throughput tRNA sequencing. Nature Methods, 12(9), 835–837. 10.1038/nmeth.3478

