## Supplemental Figures for "The Landscape of tRNA Modifications in Archaea"

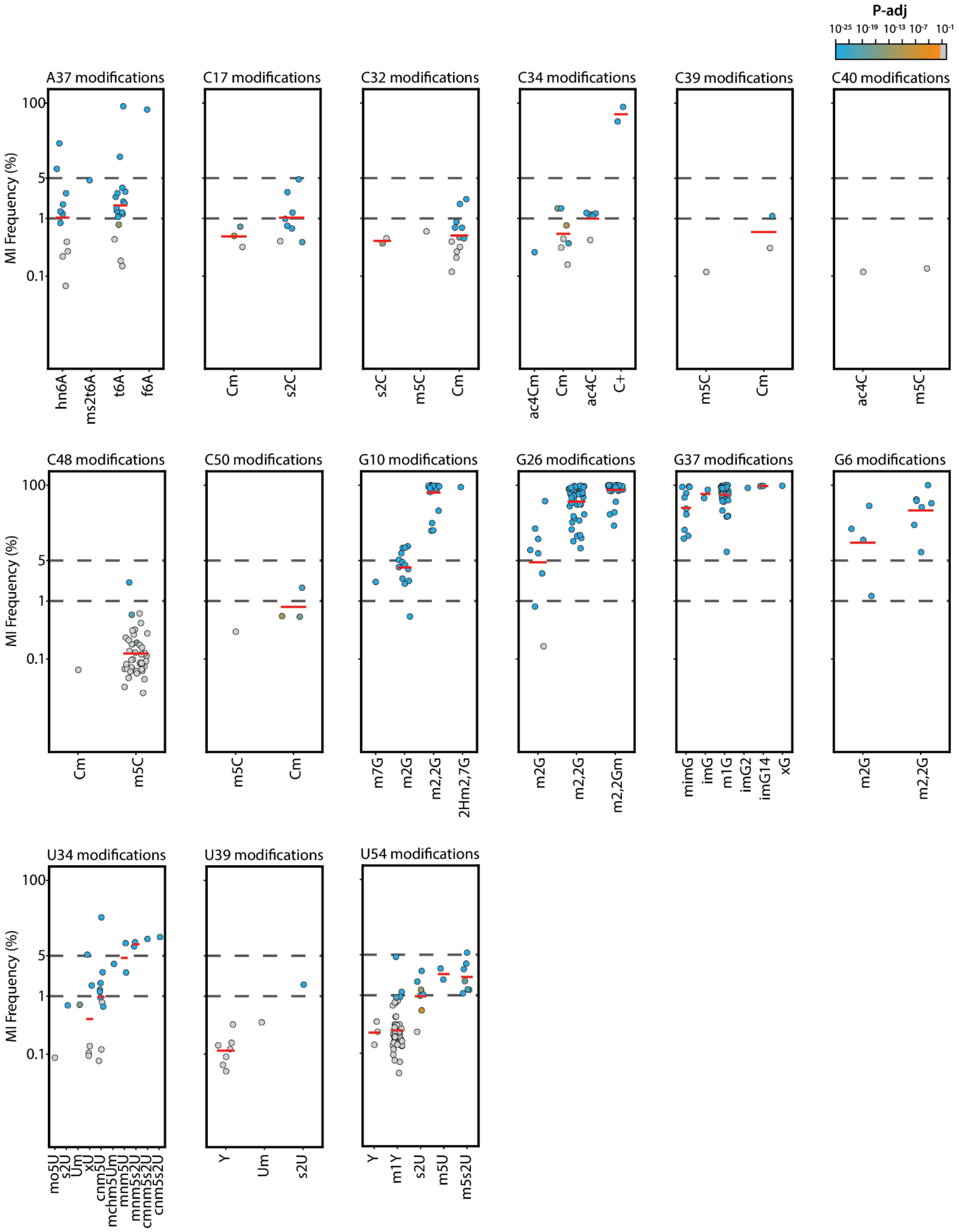

P-adj
10-25 10-19 10-13 10-7 10-1
A37 modifications
C17 modifications
C32 modifications
C34 modifications
C39 modifications
C40 modifications
C48 modifications
C50 modifications
G10 modifications
G26 modifications
G37 modifications
G6 modifications
U34 modifications
U39 modifications
U54 modifications
MI Frequency (%)
100
5
1
0.1
hn6A
ms2t6A
t6A
f6A
Cm
s2C
s2C
m5C
Cm
ac4Cm
Cm
ac4C
C+
m5C
Cm
ac4C
m5C
Cm
m5C
m5C
Cm
m7G
m2G
m2,2G
2Hm2,7G
m2G
m2,2G
m2,2Gm
mimG
imG
m1G
imG2
imG14
xG
m2G
m2,2G
mo5U
s2U
Um
xU
cnm5U
mchm5Um
mnm5U
mnm5s2U
cmnm5s2U
cnm5s2U
Y
Um
s2U
Y
m1Y
s2U
m5U
m5s2U

**Figure S1**. Misincorporation frequency (MI) is shown for a range of annotated tRNA modifications grouped by nucleotide and position (e.g., A37, C17, G10, U54). Each subplot displays individual tRNA sites as points, with color indicating the adjusted p-value (color scale at top right). Red bars denote the mean MI for the modification type. Confidence classification: 5% for A/U and 10% for G/C. Sites with MI values above these thresholds and significant adjusted p-values (p-adj < 0.05) are considered high-confidence predictions; those below the threshold but still significant are classified as moderate-confidence. Note that certain modifications, such as $m^2_2G$ display consistent distributions of MI, while others show broader variability.

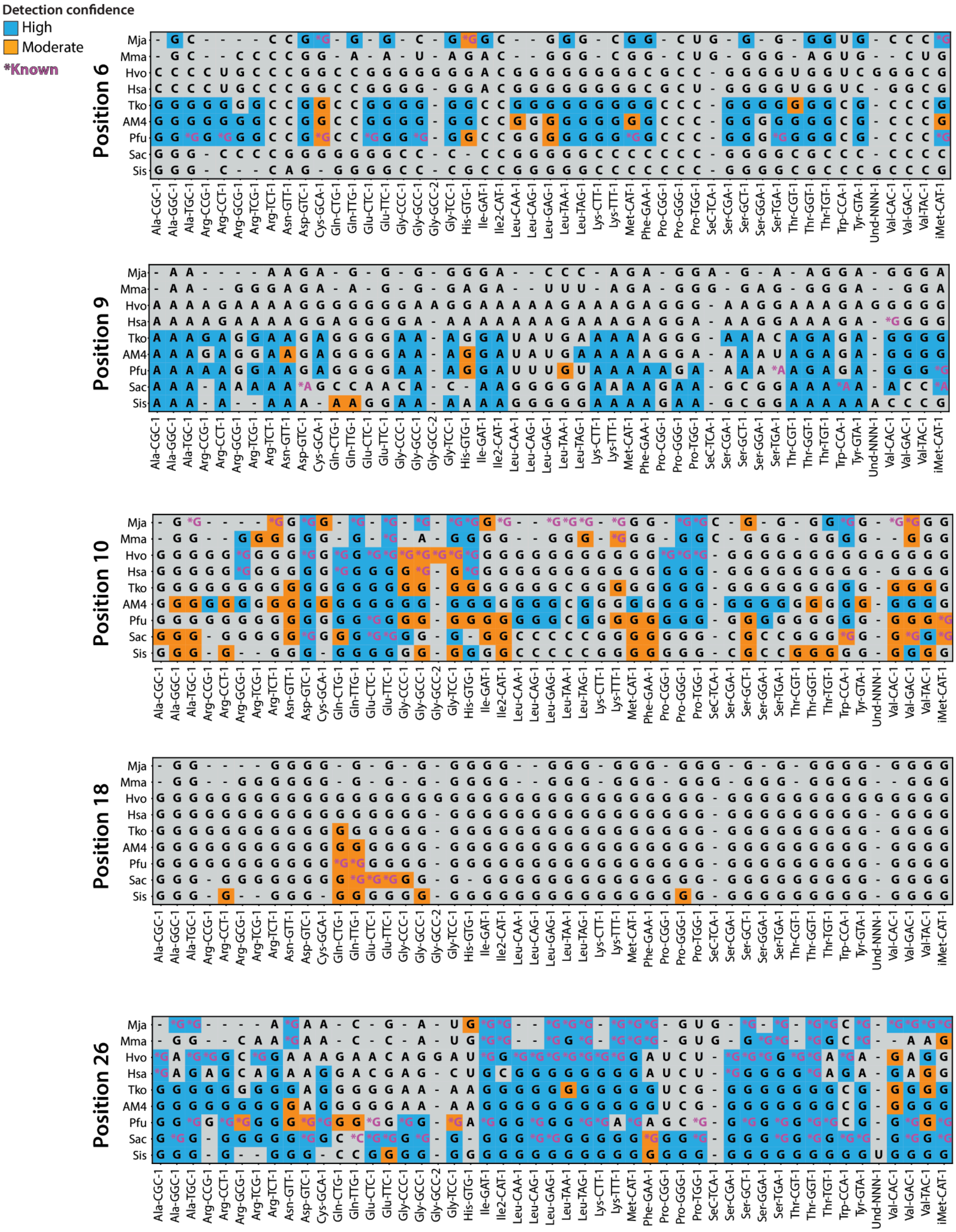
Detection confidence
High
Moderate
*Known
Position 6
Position 9
Position 10
Position 18
Position 26
Mja
Mma
Hvo
Hsa
Tko
AM4
Pfu
Sac
Sis
Ala-CGC-1
Ala-GGC-1
Ala-TGC-1
Arg-CCG-1
Arg-CCT-1
Arg-GCG-1
Arg-TCG-1
Arg-TCT-1
Asn-GTT-1
Asp-GTC-1
Cys-GCA-1
Gln-CTG-1
Gln-TTG-1
Glu-CTC-1
Glu-TTC-1
Gly-CCC-1
Gly-GCC-1
Gly-GCC-2
Gly-TCC-1
His-GTG-1
Ile-GAT-1
Ile2-CAT-1
Leu-CAA-1
Leu-CAG-1
Leu-GAG-1
Leu-TAA-1
Leu-TAG-1
Lys-CTT-1
Lys-TTT-1
Met-CAT-1
Phe-GAA-1
Pro-CGG-1
Pro-GGG-1
Pro-TGG-1
SeC-TCA-1
Ser-CGA-1
Ser-GCT-1
Ser-GGA-1
Ser-TGA-1
Thr-CGT-1
Thr-GGT-1
Thr-TGT-1
Trp-CCA-1
Tyr-GTA-1
Und-NNN-1
Val-CAC-1
Val-GAC-1
Val-TAC-1
iMet-CAT-1

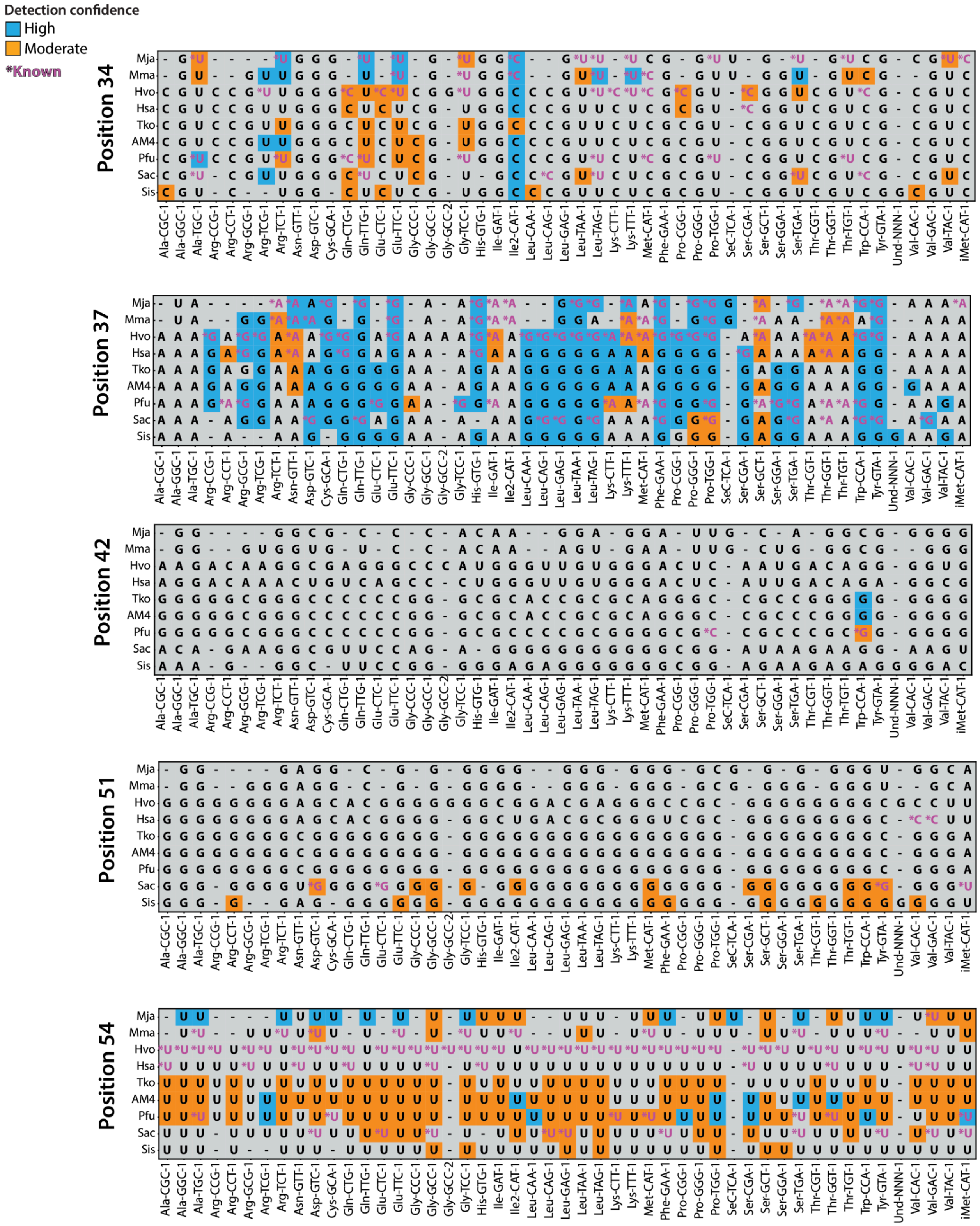

Detection confidence
High
Moderate
*Known
Position 34
Position 37
Position 42
Position 51
Position 54
Mja
Mma
Hvo
Hsa
Tko
AM4
Pfu
Sac
Sis
Ala-CGC-1
Ala-GGC-1
Ala-TGC-1
Arg-CCG-1
Arg-CCT-1
Arg-GCG-1
Arg-TCG-1
Arg-TCT-1
Asn-GTT-1
Asp-GTC-1
Cys-GCA-1
Gln-CTG-1
Gln-TTG-1
Glu-CTC-1
Glu-TTC-1
Gly-CCC-1
Gly-GCC-1
Gly-GCC-2
Gly-TCC-1
His-GTG-1
Ile-GAT-1
Ile2-CAT-1
Leu-CAA-1
Leu-CAG-1
Leu-GAG-1
Leu-TAA-1
Leu-TAG-1
Lys-CTT-1
Lys-TTT-1
Met-CAT-1
Phe-GAA-1
Pro-CGG-1
Pro-GGG-1
Pro-TGG-1
SeC-TCA-1
Ser-CGA-1
Ser-GCT-1
Ser-GGA-1
Ser-TGA-1
Thr-CGT-1
Thr-GGT-1
Thr-TGT-1
Trp-CCA-1
Tyr-GTA-1
Und-NNN-1
Val-CAC-1
Val-GAC-1
Val-TAC-1
iMet-CAT-1

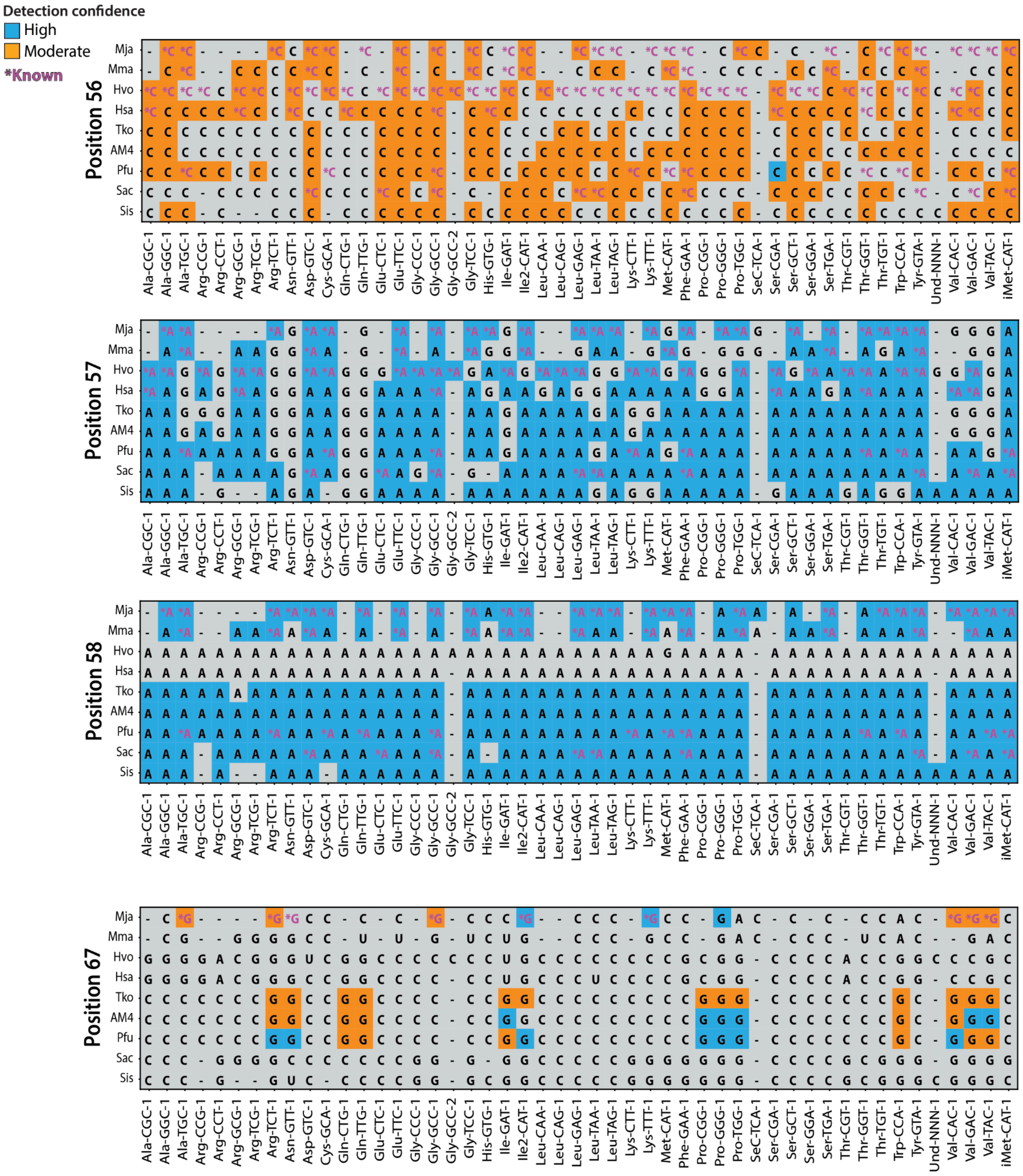
Detection confidence
High
Moderate
*Known
Position 56
Position 57
Position 58
Position 67
Mja
Mma
Hvo
Hsa
Tko
AM4
Pfu
Sac
Sis
Ala-CGC-1 Ala-GGC-1 Ala-TGC-1 Arg-CCG-1 Arg-CCT-1 Arg-GCG-1 Arg-TCG-1 Arg-TCT-1 Asn-GTT-1 Asp-GTC-1 Cys-GCA-1 Gln-CTG-1 Gln-TTG-1 Glu-CTC-1 Glu-TTC-1 Gly-CCC-1 Gly-GCC-1 Gly-GCC-2 Gly-TCC-1 His-GTG-1 Ile-GAT-1 Ile2-CAT-1 Leu-CAA-1 Leu-CAG-1 Leu-GAG-1 Leu-TAA-1 Leu-TAG-1 Lys-CTT-1 Lys-TTT-1 Met-CAT-1 Phe-GAA-1 Pro-CGG-1 Pro-GGG-1 Pro-TGG-1 SeC-TCA-1 Ser-CGA-1 Ser-GCT-1 Ser-GGA-1 Ser-TGA-1 Thr-CGT-1 Thr-GGT-1 Thr-TGT-1 Trp-CCA-1 Tyr-GTA-1 Und-NNN-1 Val-CAC-1 Val-GAC-1 Val-TAC-1 iMet-CAT-1

**Figure S2**. Predicted and known tRNA modifications across positions 6, 9, 10, 18, 26, 34, 37, 42, 51, 54, 56, 57, 58, and 67 in archaeal species, with assigned confidence levels. Each heatmap shows nucleotide identities for all annotated tRNA isotypes across archaeal species. Rows represent species (e.g., *M. jannaschii*, *T. kodakarensis*, *H. salinarum*), and columns represent individual tRNA isotypes. Colored boxes denote predicted modifications: high-confidence predictions (blue), moderate-confidence predictions (orange), and known modifications (purple asterisks). Sites without significant misincorporation signals are shown in gray. Positions with '-' indicate a missing tRNA that is not present in a specific tRNA set, or a tRNA that was not captured by sequencing. This comparative map provides a detailed view of how specific modifications are deployed across phylogeny and tRNA substrates, supporting broader conclusions of clade- and identity-specific modification predictions. However, it's important to note there are limitations in RT-based detection of non-Watson-Crick disrupting modifications, such as the "invisible" $m^5U54$, which are not observed despite being "known" in the reference set. A derivative of $m^5U54$, that is $m^5s^2U54$, results in low to moderately detectable misincorporations but should still be considered with caution when drawing comparisons and inferring patterns of variation. Similarly, Cm56 does not directly perturb the RT but occasionally has a subtle impact via a sugar pucker of the 3' carbon on the ribose that is positioned above the plane. This explains the observation made for many Nm modifications and their low to moderately detectable misincorporations.

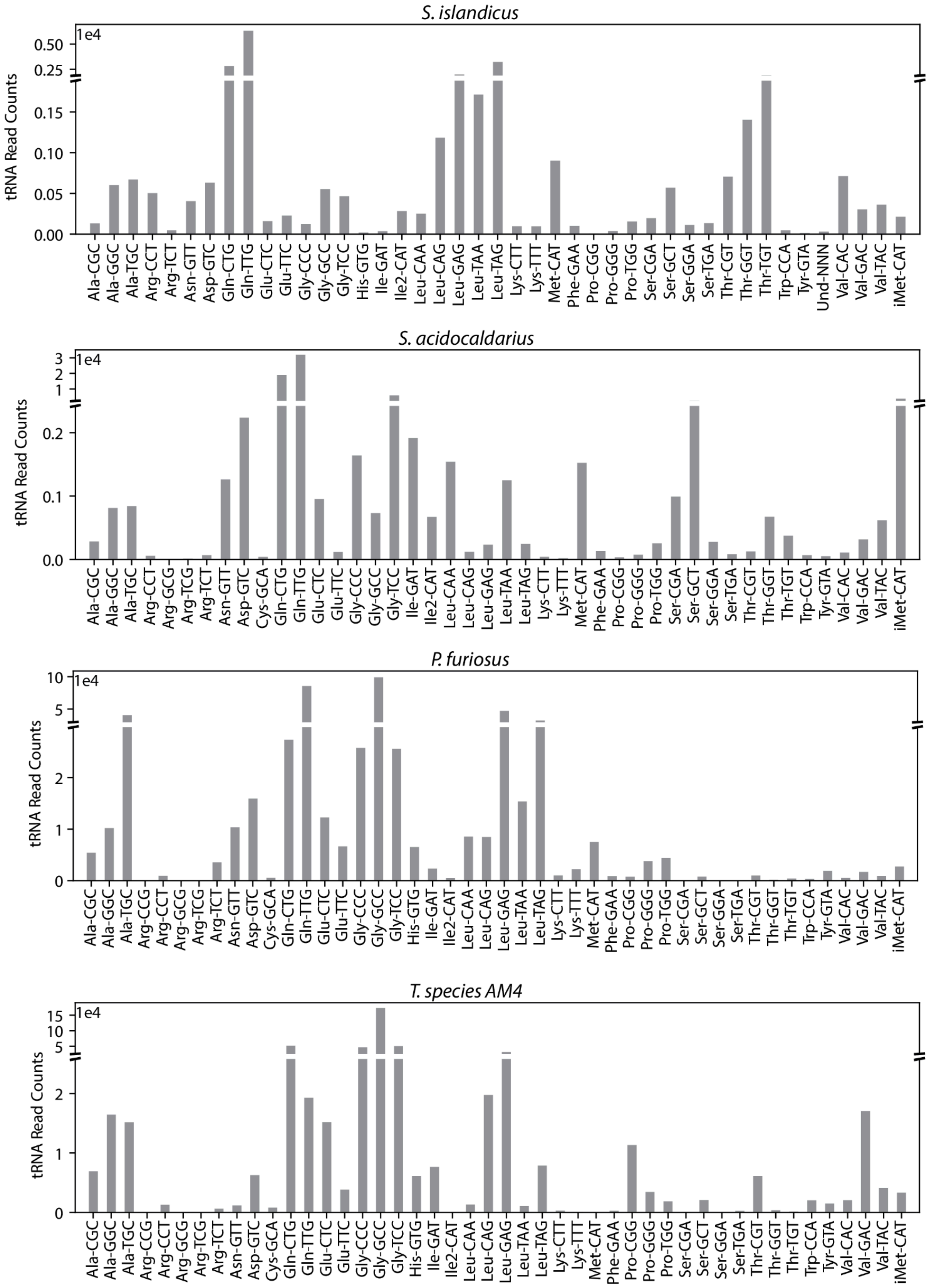

S. islandicus
1e4
0.50
0.25
0.15
0.10
0.05
0.00
tRNA Read Counts
Ala-CGC Ala-GGC Ala-TGC Arg-CCT Arg-TCT Asn-GTT Asp-GTC Gln-CTG Gln-TTG Glu-CTC Glu-TTC Gly-CCC Gly-GCC Gly-TCC His-GTG Ile-GAT Ile2-CAT Leu-CAA Leu-CAG Leu-GAG Leu-TAA Leu-TAG Lys-CTT Lys-TTT Met-CAT Phe-GAA Pro-CGG Pro-GGG Pro-TGG Ser-CGA Ser-GCT Ser-GGA Ser-TGA Thr-CGT Thr-GGT Thr-TGT Trp-CCA Tyr-GTA Und-NNN Val-CAC Val-GAC Val-TAC iMet-CAT
S. acidocaldarius
1e4
3
2
1
0.2
0.1
0.0
tRNA Read Counts
Ala-CGC Ala-GGC Ala-TGC Arg-CCT Arg-GCG Arg-TCG Arg-TCT Asn-GTT Asp-GTC Cys-GCA Gln-CTG Gln-TTG Glu-CTC Glu-TTC Gly-CCC Gly-GCC Gly-TCC Ile-GAT Ile2-CAT Leu-CAA Leu-CAG Leu-GAG Leu-TAA Leu-TAG Lys-CTT Lys-TTT Met-CAT Phe-GAA Pro-CGG Pro-GGG Pro-TGG Ser-CGA Ser-GCT Ser-GGA Ser-TGA Thr-CGT Thr-GGT Thr-TGT Trp-CCA Tyr-GTA Val-CAC Val-GAC Val-TAC iMet-CAT
P. furiosus
1e4
10
5
2
1
0
tRNA Read Counts
Ala-CGC Ala-GGC Ala-TGC Arg-CCG Arg-CCT Arg-GCG Arg-TCG Arg-TCT Asn-GTT Asp-GTC Cys-GCA Gln-CTG Gln-TTG Glu-CTC Glu-TTC Gly-CCC Gly-GCC Gly-TCC His-GTG Ile-GAT Ile2-CAT Leu-CAA Leu-CAG Leu-GAG Leu-TAA Leu-TAG Lys-CTT Lys-TTT Met-CAT Phe-GAA Pro-CGG Pro-GGG Pro-TGG Ser-CGA Ser-GCT Ser-GGA Ser-TGA Thr-CGT Thr-GGT Thr-TGT Trp-CCA Tyr-GTA Val-CAC Val-GAC Val-TAC iMet-CAT
T. species AM4
1e4
15
10
5
2
1
0
tRNA Read Counts
Ala-CGC Ala-GGC Ala-TGC Arg-CCG Arg-CCT Arg-GCG Arg-TCG Arg-TCT Asn-GTT Asp-GTC Cys-GCA Gln-CTG Gln-TTG Glu-CTC Glu-TTC Gly-CCC Gly-GCC Gly-TCC His-GTG Ile-GAT Ile2-CAT Leu-CAA Leu-CAG Leu-GAG Leu-TAA Leu-TAG Lys-CTT Lys-TTT Met-CAT Phe-GAA Pro-CGG Pro-GGG Pro-TGG Ser-CGA Ser-GCT Ser-GGA Ser-TGA Thr-CGT Thr-GGT Thr-TGT Trp-CCA Tyr-GTA Val-CAC Val-GAC Val-TAC iMet-CAT

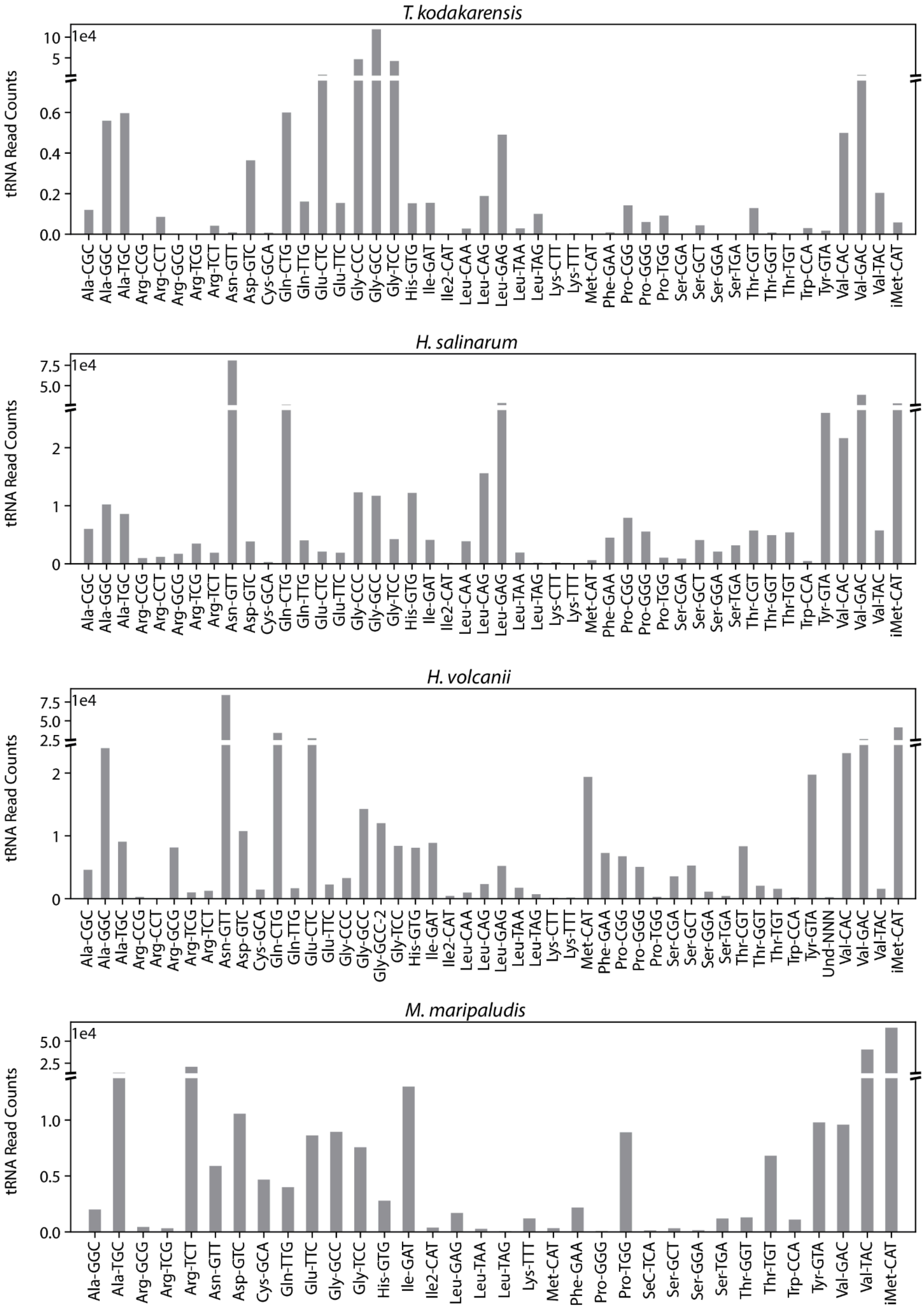

T. kodakarensis
H. salinarum
H. volcanii
M. maripaludis
tRNA Read Counts
1e4
Ala-CGC
Ala-GGC
Ala-TGC
Arg-CCG
Arg-CCT
Arg-GCG
Arg-TCG
Arg-TCT
Asn-GTT
Asp-GTC
Cys-GCA
Gln-CTG
Gln-TTG
Glu-CTC
Glu-TTC
Gly-CCC
Gly-GCC
Gly-GCC-2
Gly-TCC
His-GTG
Ile-GAT
Ile2-CAT
Leu-CAA
Leu-CAG
Leu-GAG
Leu-TAA
Leu-TAG
Lys-CTT
Lys-TTT
Met-CAT
Phe-GAA
Pro-CGG
Pro-GGG
Pro-TGG
SeC-TCA
Ser-CGA
Ser-GCT
Ser-GGA
Ser-TGA
Thr-CGT
Thr-GGT
Thr-TGT
Trp-CCA
Tyr-GTA
Und-NNN
Val-CAC
Val-GAC
Val-TAC
iMet-CAT

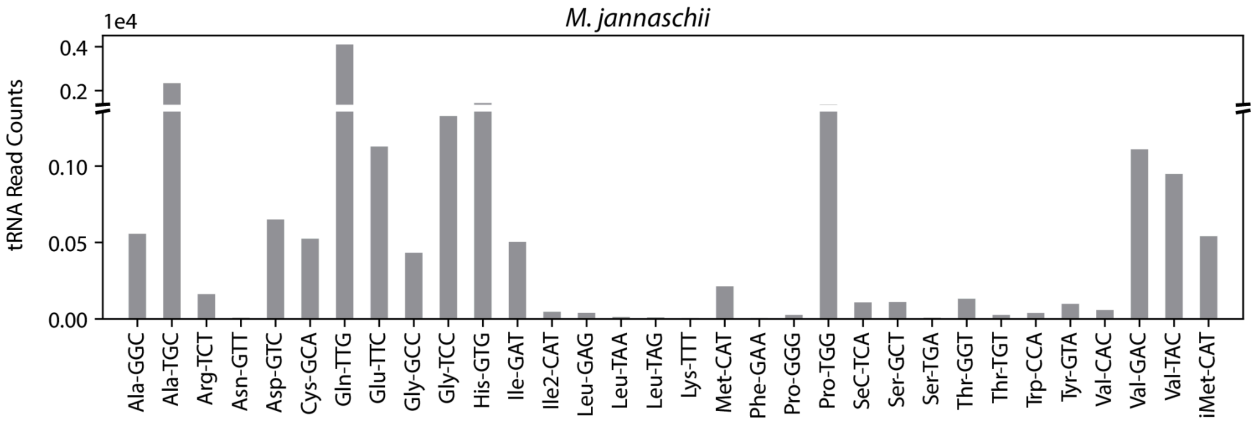


**Figure S3**. Total tRNA read counts mapped for each tRNA across all archaeal species. Bar plots show the number of reads mapped to each species. Read counts are shown on the y-axis (log scale), and tRNAs are ordered by amino acid and anticodon. These data reflect the sequencing depth and tRNA abundance for each species. Read coverage was robust across most species. Notably, *S. islandicus* and *M. jannaschii* had comparatively low overall tRNA read depth. However, for tRNAs that were captured, misincorporation profiles are independent of coverage.

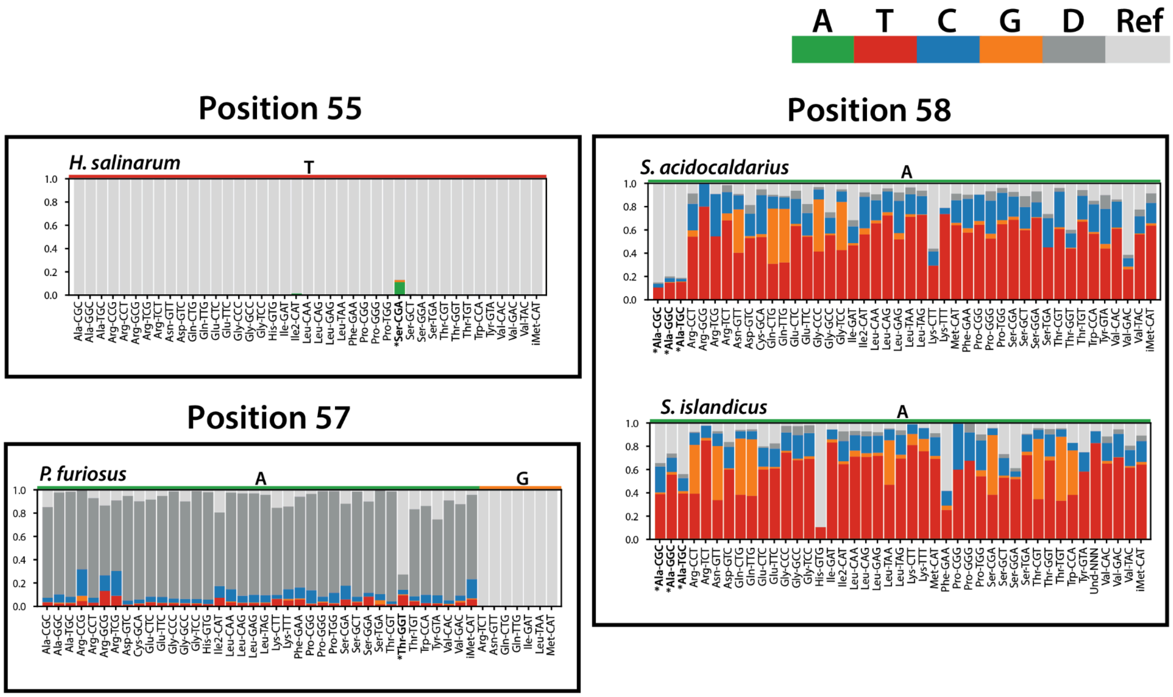


**Figure S4**. Outliers (marked with *) for Positions 55, 57, and 58, as referenced in Figure 1C. Pseudouridine at position 55 was consistently undetectable across species, as expected given its minimal impact on reverse transcription. However, in *H. salinarum* $\text{Ser}^{\text{CGA}}$, we observed an unusually high misincorporation rate and a significant p-value at this position. While pseudouridine alone is unlikely to account for this signal, it may reflect the presence of a related or modified derivative, such as $m^1Y$, which has been reported at nearby positions, like 54, but may occur at 55 in this specific context. $m^1I$ at position 57 was consistently detectable, however in *P.furiosus* $\text{Thr}^{\text{GGU}}$, we observed an unusually low misincorporation rate that suggests only partial modification. G occurs at this position for 7 tRNAs (Arg-TCT, Asn-GTT, etc), showing a lack of MI, for comparison. Similarly, $m^1A$ at position 58 was consistently detectable; however, for the Thermoprotei *S. acidocaldarius* and *S. islandicus* $\text{tRNA}^{\text{Ala}}$, we observed unusually low misincorporation rates, suggesting partial modification or tRNA-specific regulation. There are additional examples of tRNAs with unexpectedly low misincorporation rates such as His-GTG, where the read depth is low which results in reduced rates of misincorporation and lack of expected modification patterns associated with $m^1A58$. In addition, for tRNAs with no read read counts the misincorporation is not displayed.

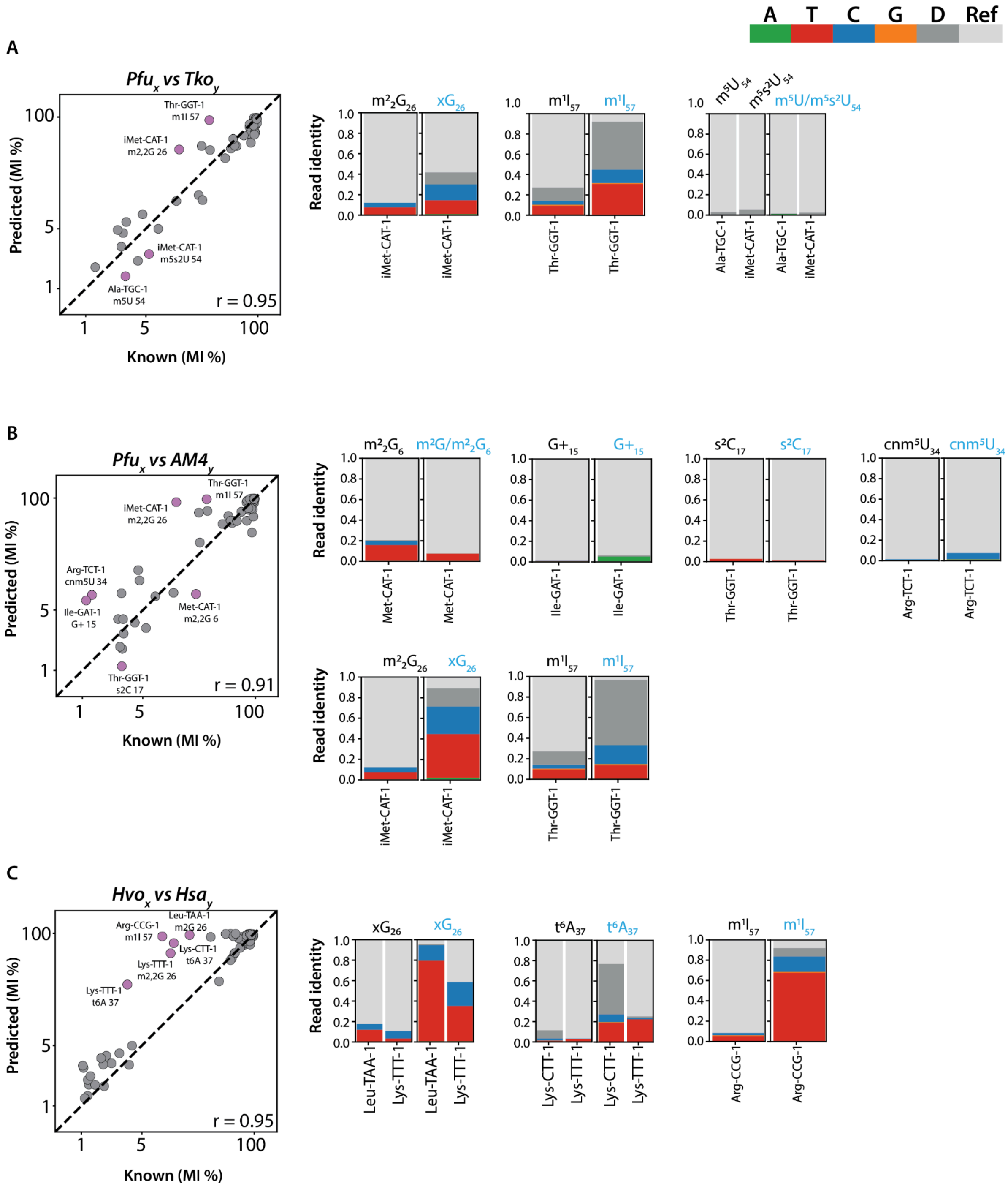
A T C G D Ref
A
Pfu_x vs Tko_y
Thr-GGT-1 m1I 57
iMet-CAT-1 m2,2G 26
iMet-CAT-1 m5s2U 54
Ala-TGC-1 m5U 54
r = 0.95
Predicted (MI %)
Known (MI %)
Read identity
m²₂G₂₆ xG₂₆
m¹I₅₇ m¹I₅₇
m⁵U₅₄ m⁵s²U₅₄ m⁵U/m⁵s²U₅₄
iMet-CAT-1
Thr-GGT-1
Ala-TGC-1
B
Pfu_x vs AM4_y
Thr-GGT-1 m1I 57
iMet-CAT-1 m2,2G 26
Arg-TCT-1 cnm5U 34
Ile-GAT-1 G+ 15
Met-CAT-1 m2,2G 6
Thr-GGT-1 s2C 17
r = 0.91
m²₂G₆ m²G/m²₂G₆
G+₁₅ G+₁₅
s²C₁₇ s²C₁₇
cnm⁵U₃₄ cnm⁵U₃₄
Met-CAT-1
Ile-GAT-1
Thr-GGT-1
Arg-TCT-1
m²₂G₂₆ xG₂₆
m¹I₅₇ m¹I₅₇
iMet-CAT-1
Thr-GGT-1
C
Hvo_x vs Hsa_y
Arg-CCG-1 m1I 57
Leu-TAA-1 m2G 26
Lys-CTT-1 t6A 37
Lys-TTT-1 m2,2G 26
Lys-TTT-1 t6A 37
r = 0.95
xG₂₆ xG₂₆
t⁶A₃₇ t⁶A₃₇
m¹I₅₇ m¹I₅₇
Leu-TAA-1
Lys-TTT-1
Lys-CTT-1
Arg-CCG-1

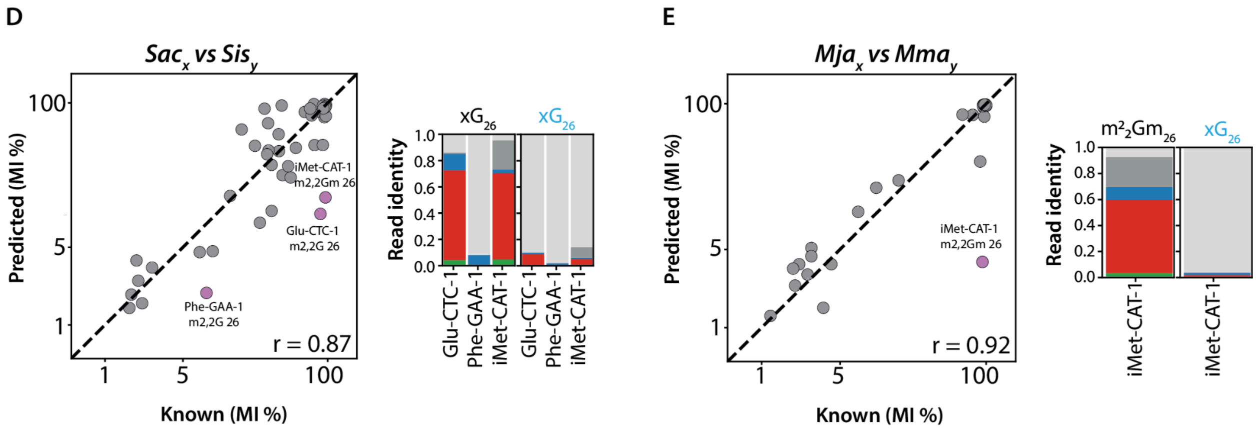


**Figure S5**. Overview of homology-based tRNA modifications predictions. A-E) Pearson correlation between known modifications vs homology-predicted modifications. Each point in the scatter plot corresponds to a specific modification type and site from homologous tRNAs. The diagonal line represents perfect correlation (r = 1) between known and predicted MI. Outliers (± 2 SD) are highlighted (purple) and annotated with their corresponding tRNA, modification, and position. The read identity (pattern of misincorporation) for select outliers is depicted next to the scatter plot, with known modification position (black) compared to newly predicted (blue) (See all outliers in Supplemental Figures). A) *P. furiosus (Pfu)* vs *T. kodakarensis (Tko)*. Select outliers iMet$^{CAU}$ $m^2_2G$ at position 26 and Thr$^{GGU}$ $m^1I$ at position 57. B) *P. furiosus (Pfu)* vs *T. species* AM4 *(*AM4*)*. Select outliers Met$^{CAU}$ $m^2_2G$ at position 6, and Arg$^{UCU}$ cnm$^5$U at position 34. C) *H. volcanii (Hvo)* vs *H. salinarum (Hsa)*. Select outliers Leu$^{UAA}$ $m^2G$, Lys$^{UUU}$ $m^2_2G$ at position 26, and Lys$^{CUU}$, Lys$^{UUU}$ $t^6A$ at position 37. D) *S. acidocaldarius (Sac) vs S. islandicus (Sis)*. All outliers Glu$^{CUC}$ and Phe$^{GAA}$ $m^2_2G$, and iMet$^{CAU}$ $m^2_2Gm$ at position 26. E) *M. jannaschi (Mja)* vs *M. maripaludis (Mma)*. Outlier iMet$^{CAU}$ $m^2_2Gm$ at position 26. For positions with more than one modification type (e.g., $m^2G/m^2_2G_{26}$), "x" was used. This notation is also used for most predictions, as exact modification types weren't ascertained.

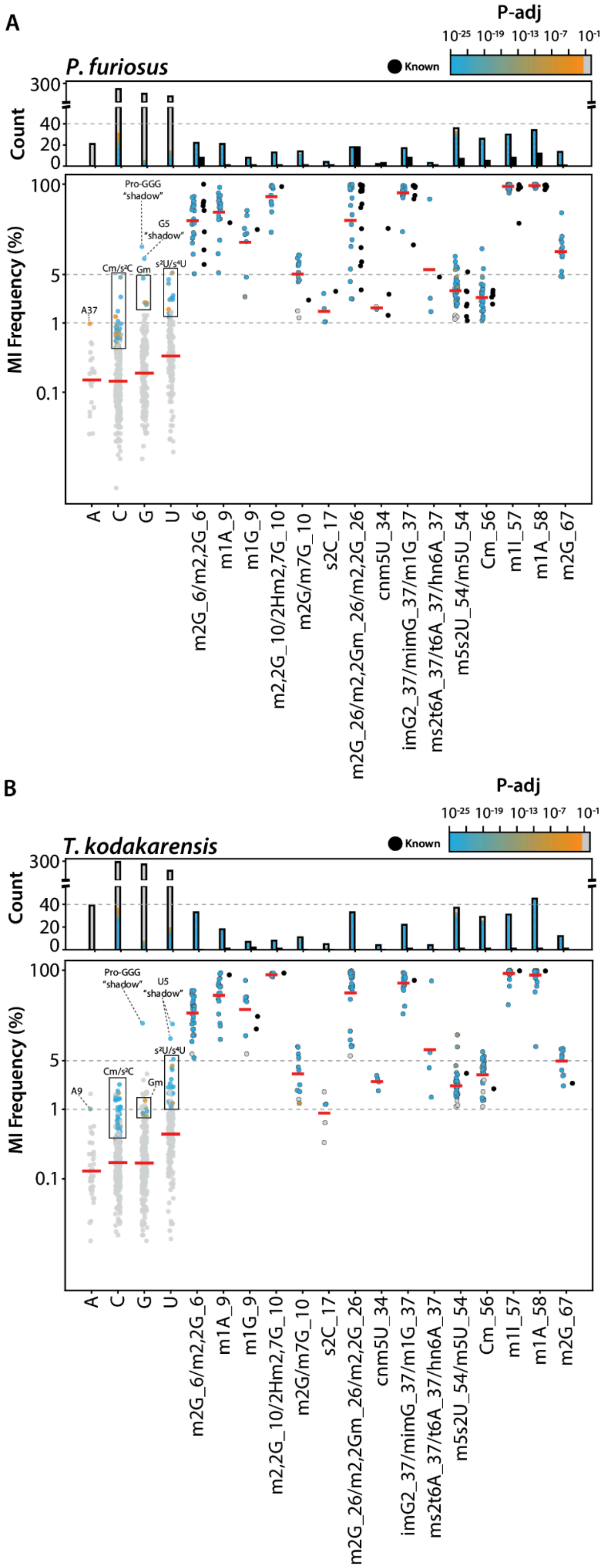
A
P-adj
10-25 10-19 10-13 10-7 10-1
Known
P. furiosus
Count
300
40
20
0
MI Frequency (%)
100
5
1
0.1
Pro-GGG "shadow"
G5 "shadow"
Cm/s²C
Gm
s²U/s⁴U
A37
A
C
G
U
m2G_6/m2,2G_6
m1A_9
m1G_9
m2,2G_10/2Hm2,7G_10
m2G/m7G_10
s2C_17
m2G_26/m2,2Gm_26/m2,2G_26
cnm5U_34
imG2_37/mimG_37/m1G_37
ms2t6A_37/t6A_37/hn6A_37
m5s2U_54/m5U_54
Cm_56
m1I_57
m1A_58
m2G_67
B
P-adj
10-25 10-19 10-13 10-7 10-1
Known
T. kodakarensis
Count
300
40
20
0
MI Frequency (%)
100
5
1
0.1
Pro-GGG "shadow"
U5 "shadow"
s²U/s⁴U
Cm/s²C
Gm
A9
A
C
G
U
m2G_6/m2,2G_6
m1A_9
m1G_9
m2,2G_10/2Hm2,7G_10
m2G/m7G_10
s2C_17
m2G_26/m2,2Gm_26/m2,2G_26
cnm5U_34
imG2_37/mimG_37/m1G_37
ms2t6A_37/t6A_37/hn6A_37
m5s2U_54/m5U_54
Cm_56
m1I_57
m1A_58
m2G_67

**Figure S6**. Known archaeal tRNA modifications and their misincorporation (MI) rates compared to predicted sites of modification for *P. furiosus* and *T. kodakarensis.* A) Misincorporation (MI) frequencies of known (black points) and newly predicted (blue-orange gradient) tRNA modifications in *P. furiosus*. The x-axis lists individual modification types, and the y-axis shows their observed or predicted MI frequency. The bar plot at the top displays the total counts for each category, with color gradients corresponding to adjusted p-values. B) Equivalent MI frequency plot for *T. kodakarensis*, again showing known (black) and predicted (blue-orange) tRNA modifications, along with total counts in the bar plot above. False-positive predictions are highlighted along with their corresponding explanations. In brief, these are composed of unannotated modifications and non-random sequencing artifacts. Generally, the misincorporation frequencies of predicted modifications closely matched those of experimentally validated sites. However, several notable exceptions ("false positives") emerged. Specifically, multiple cytosine positions exhibited significant adjusted p-values, despite lacking previous annotations. For example, position C17, a site known to be modified with either 2'-O-methylcytidine (Cm) or 2-thiocytidine ($s^2C$), was not significantly annotated with Cm17 in any clade-specific reference species, resulting in these modifications at analogous positions (such as C6, C17, and C42) in *P. furiosus* and *T. kodakarensis* being called as false positives. Similarly, guanine and uridine-derived false positives are called specifically at positions known to be modified (U8, U39, G18, and G51) with 4-thiouridine ($s^4U$), 2-thiouridine ($s^2U$), and 2'-O-methylguanine (Gm). These signals likely represent genuine modifications that escaped homologous-prediction annotations rather than true false positives, and suggest the possible presence of some previously undetected modifications within these species. Lastly, there are occasional false positives that occur as a consequence of upstream modifications followed by short repetitive nucleotide sequences that we have termed "shadow" misincorporations. For example, G34 of $Pro^{GGG}$ is the most commonly seen shadow across all species. Similarly, some U5 of $tRNA^{Gly}$ and G5 of $tRNA^{Thr}$ are also observed to present these shadows.

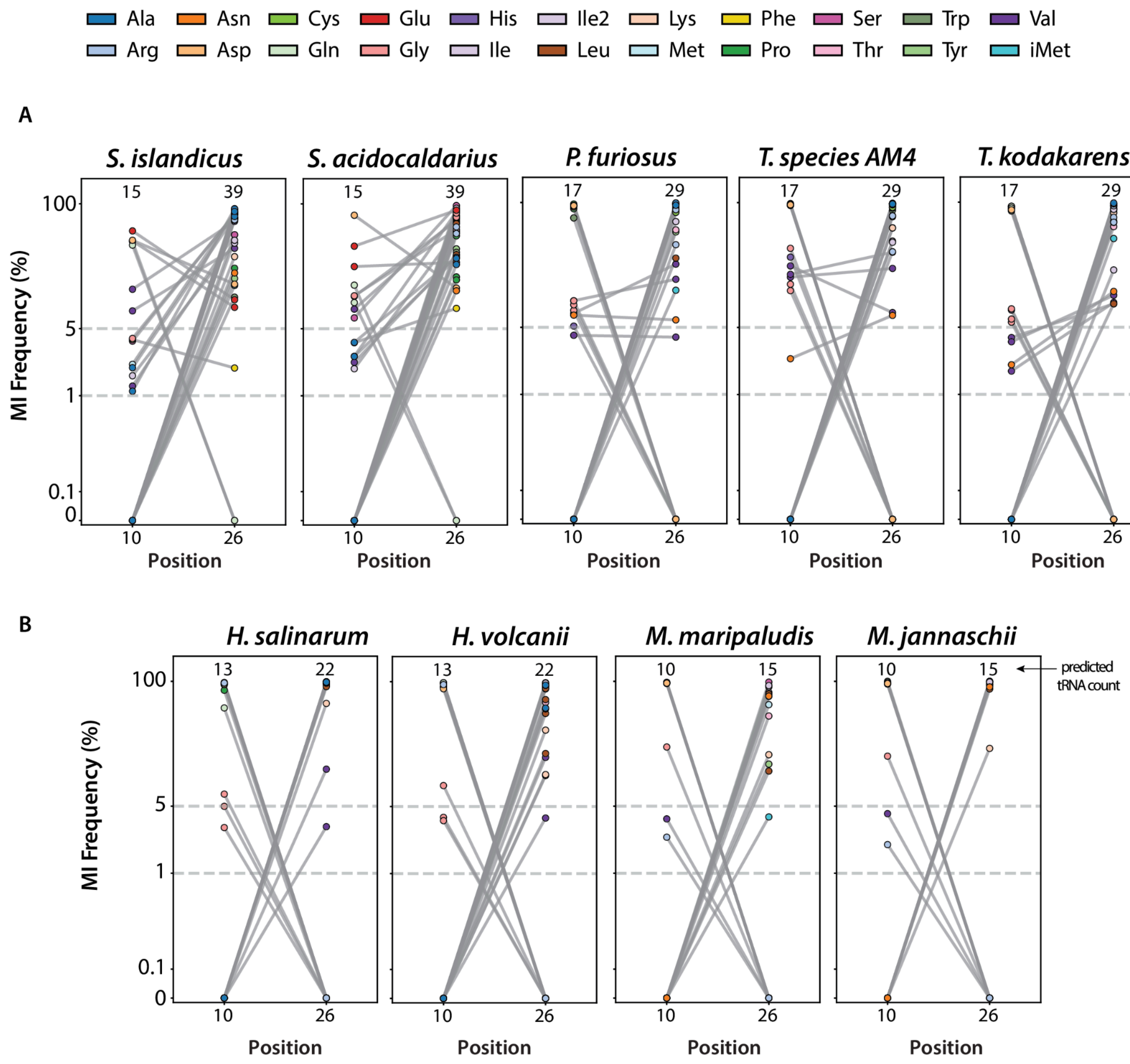


**Figure S7**. Coordinated modification pattern of modification in positions 10 and 26 across archaeal tRNAs. Paired misincorporation frequencies are shown for individual tRNAs predicted to be modified at position 10 and/or position 26 across all species. A) Thermococci and Thermoprotei species (*S. islandicus, S. acidocaldarius, P. furiosus, T. species* AM4, *T. kodakarensis*) exhibit co-occurrence of modifications at both positions within the same tRNA. This is especially prominent in Thermoprotei, where high-confidence predictions indicate that many tRNAs, such as $tRNA^{Asp}$, $tRNA^{Glu}$, and $tRNA^{Val}$, carry both $m^2_2G10$ and $m^2_2G26$. In contrast, Thermococci species display a similar trend, but with most misincorporations occurring at position 10, suggesting that $m^2G10$ is paired with $m^2_2G26$ in the same tRNA. B) In Halobacteria (*H. salinarum, H. volcanii*) and Methanococci (*M. maripaludis, M. jannaschii*), positions 10 and 26 are more often modified in mutually exclusive subsets of tRNAs, suggesting distinct substrate preferences or competitive enzyme activity. Each line represents one tRNA, connecting MI frequencies at position 10 (left) and 26 (right), with point color indicating isotype identity. Numbers above each panel reflect the number of predicted modifications at the position. Supporting data for these plots is available in Table S6.

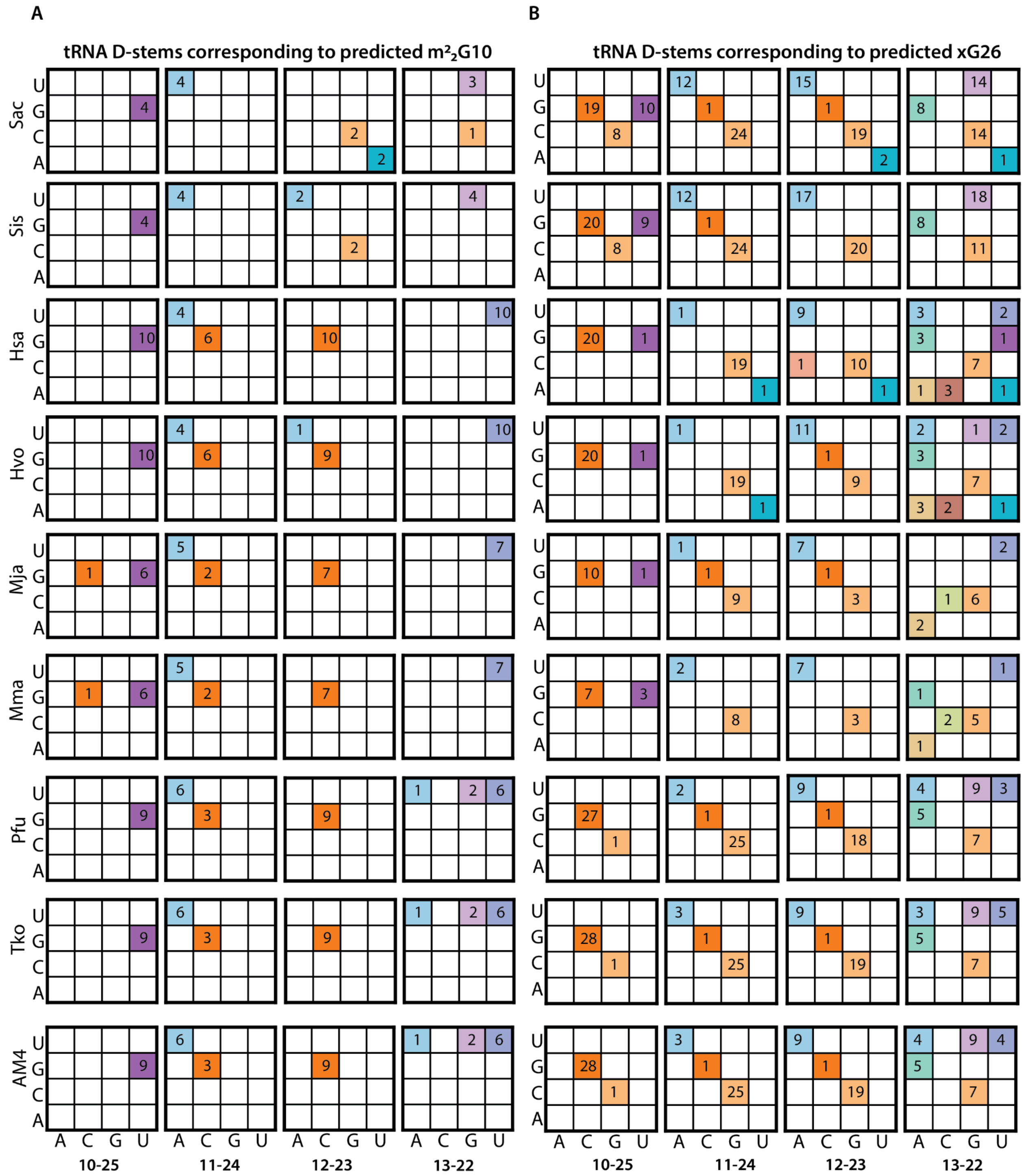


**Figure S8**. Conserved D-stem pairing patterns associated with predicted $m^2_2$G10 and xG26 modifications across archaeal tRNAs. Each map illustrates the frequency of specific D-stem base pairings in tRNAs predicted to be modified as position 10 (panel A) or position 26 (panel B), across nine representative archaeal species. Counts represent the number of predicted modified tRNAs that contain the indicated nucleotide pairs at canonical D-stem positions: 10-25, 11-24,

12-23, and 13-22. A) tRNAs predicted to carry $m^2_2G10$ are strongly associated with G10oU25 base pair, a known identity element for Trm11-mediated modification. Additional motifs such as U13oU22 and U13oA22 are frequently observed in Euryarchaeota species (e.g., *H. volcanii, M. jannaschii*) and may contribute to substrate specificity. In contrast, some Thermoprotei species (e.g., *S. acidocaldarius*, *S. islandicus*) exhibit $m^2_2G10$ modifications in tRNA that lack these canonical features, suggesting relaxed or alternative recognition in this clade. B) xG26-modified tRNA ($m^2G$, $m^2_2G$, $m^2_2Gm$) show a broader set of D-stem pairing preferences. While canonical G10oU25 pairs are common in many species, they tend to exclude position 26 modification in Euryarchaeota, consistent with prior findings in *P. furiosus*. This exclusionary relationship is not maintained in Thermoprotei, which frequently exhibit co-modification at both positions and show a preference for alternative motifs, such as C12=G23 or A12oU23. These data emphasize the strong evolutionary conservation and divergence of D-arm identity elements that guide enzyme-substrate recognition. Patterns suggest that shared structural features enable coordinated modification at positions 10 and 26 in some lineages, while other species may exhibit unique targeting.

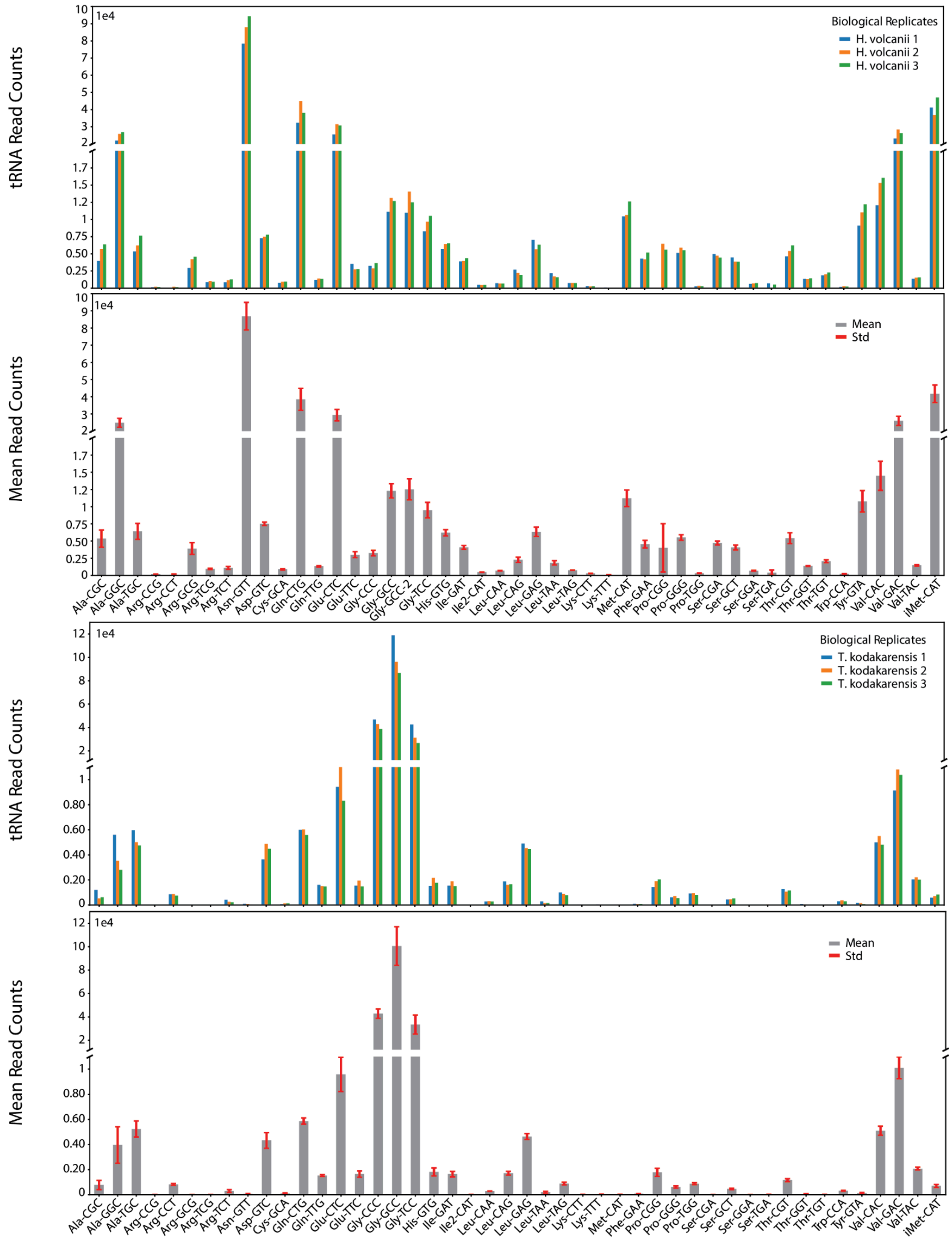


**Figure S9**. Total tRNA read counts across three biological replicates for *H. volcanii* and *T. kodakarensis*. Two pairs of panels for each species. Top panel includes reads from each replicate, bottom panel is a summary of each, with mean and standard deviation (std).
